# Colony geometric complexity predicts the co-occurrence of coral photosymbiont genera

**DOI:** 10.64898/2026.09.30.755836

**Authors:** Alyssa M. Tomkies, Evan M. Heit, Teina Rongo, Michael Kühl, David J. Suggett, Simon K. Davy, Matthew R. Nitschke

## Abstract

The endosymbiosis between tropical reef-building corals and dinoflagellate algae (family Symbiodiniaceae) underpins the healthy functioning of coral reef ecosystems. Although dominant host-symbiont pairings differ across spatial scales (e.g., bioregion, reef zone, and depth), corals often associate with multiple symbionts. Where diverse symbionts occur together, their local-scale ecology is poorly understood. Moreover, whilst coral morphological plasticity generates a structurally complex spatial habitat for endosymbiosis, the role of host geometry in facilitating symbiont co-occurrence is completely unexplored. Here, we integrated high-throughput sequencing of the ITS2 gene region with digital 3D photogrammetry to evaluate symbiont distribution within coral structural habitat and determine the influence of host morphology on symbiont community composition. Symbiont composition was characterised along a vertical gradient (coral apex to base) across 50 *Hydnophora microconos* colonies from Rarotonga, Cook Islands. We identified dominance by the genera *Durusdinium* and *Cladocopium*, with inter-genus co-occurrence recognised in 60% of coral colonies. The co-occurrence of two *Cladocopium* lineages (C3f and C3k) was validated based on congruence with the plastid minicircle psbA non-coding region (psbA^ncr^). Geometric quantification from digital 3D reconstructions of coral colonies revealed that single symbiont taxon colonies were morphologically distinct, whilst multi-symbiont colonies exhibited a broader morphological range. Where *Cladocopium*-*Durusdinium* co-occurrence was identified, surface orientation indicated increased *Durusdinium* prevalence toward the base of coral colonies, corresponding to regions with the highest occlusion of modelled downwelling light. Collectively, these findings suggest that coral structural heterogeneity supports Symbiodiniaceae diversity and highlights the value of resolving microhabitat-level variation to understand symbiont co-occurrence.

## Introduction

Photosynthetic endosymbionts (family Symbiodiniaceae, hereafter ‘symbionts’) are integral to scleractinian coral function and, thus, the productivity of tropical coral reef ecosystems. Through the translocation of carbon-based photosynthates, algal symbionts can supply more than 100% of host energetic requirements, underpinning coral performance in otherwise nutrient poor waters [1,2]. The stability of this partnership, and consequently the persistence of coral reefs, is vulnerable to environmental perturbations [3,4], where elevated thermal conditions in particular drive symbiotic breakdown, causing host bleaching and heightened mortality risk [5,6]. Yet, the Symbiodiniaceae exhibit substantial genetic and functional variation across 16 genus-equivalent lineages [7], with host fitness influenced by symbiont physiology [8–10]. Accordingly, association with functionally distinct symbionts can modulate thermal tolerance and bleaching susceptibility, supporting host coral resilience to environmental stress [11–15].

Across multiple spatial scales, host-symbiont compatibility and environmental variation shape symbiont distributions [16,17]. In the highly speciose genus *Cladocopium*, widespread symbiotic taxa occupy diverse irradiances and thermal environments, with particular prevalence in the Indo-Pacific [18,19,7,20]. Members of the genus *Durusdinium*, also common in the Indo-Pacific, are more frequently associated with warmer or highly turbid environments [16], suggesting increased resilience to thermal extremities [21]. At reduced levels of spatial hierarchy, patterns in symbiont distribution remain evident. Within the same host species, compatible symbiont associations can differ across local reefs or from inshore to offshore areas [22,23] in correlation with local variation in abiotic conditions [24,25]. Similarly, symbiont distributions vary along depth gradients, where changing irradiance and temperature alter the environmental suitability of host-symbiont associations [26–29].

Environmental heterogeneity is not limited to broad spatial scales, also occurring at the scale of individual coral colonies where resource variation generates diverse microenvironments across the three-dimensional colony surface [30,31]. Reef-building scleractinian corals are typically colonial organisms, forming modular structures in which genetically identical units (polyps) replicate to drive skeletal modification [32,33]. Unlike the fixed geometry of unitary organisms, this modularity facilitates growth pattern plasticity [34,35], creating complexity within species-specific bounds of structural architecture. Across ecological systems, structural complexity enhances microhabitat diversity, influencing biodiversity and aiding the partitioning of functionally distinct organisms across niche space [36–38]. The influence of coral colony heterogeneity on symbiont distribution, however, remains poorly characterised. Growing evidence indicates that many coral taxa have the capacity to associate with multiple Symbiodiniaceae [39,40], with a recent assessment detecting 9.6% global *Cladocopium*-*Durusdinium* co-occurrence as a conservative estimate that was substantially elevated through intra-colony sampling and high-throughput genotyping methods [41]. Variation in symbiont taxa has been identified along vertically oriented colony surfaces, where downwelling irradiance has been implicated in intra-colony distribution, yet these patterns are not ubiquitous [42–45], suggesting a complex interplay of ecological mechanisms yet to be quantified. Whether within colony morphological (‘structural habitat’) differentiation shapes symbiont communities at the microscale, thereby contributing to niche partitioning, remains unresolved; such variation is plausible yet may have been obscured by sampling resolution coarser than the microscale at which it operates.

Recent evaluations of the structural shape of reef-building corals utilizing continuous, geometry-specific variables have advanced the characterisation of coral morphology [46–48], building on traditional description according to broad growth forms (e.g., massive, arborescent, digitate, encrusting). The increasing application of photogrammetry as a digital tool has supported this advancement [49]; in particular, applying digital photogrammetric techniques at the scale of individual colonies has enabled comprehensive evaluation of structural heterogeneity [50–52]. Such methods facilitate a high-resolution approach to evaluating colony geometry and quantifying structural microenvironments together with symbiont composition.

Here, we aim to evaluate the hypothesis that symbiont co-occurrence is predicted by coral colony morphology. To address this, we integrate digital photogrammetry and high-throughput sequencing to determine the distribution of symbionts within coral habitat space. We determined the structural geometry of coral colonies and quantified patterns of symbiont composition in relation to morphological variation. Subsequently, we assessed the three-dimensional spatial position of tissue samples in relation to modelled ambient occlusion as a driver of multi-symbiont assemblages.

## Methods

### Species selection and sample collection

All samples were collected from the coral species *Hydnophora microconos* [53]*. H. microconos* has a broad geographic distribution, spans growth forms from massive to encrusting, and is readily identified by conical skeletal projections (monticules) [54–56]. Symbiont associations in *H. microconos* encompass distinct intra- and inter-genus taxa [45], and hence make it a strong candidate species for investigating the role of host colony geometry in symbiont co-occurrence.

Fifty *H. microconos* colonies were identified in shallow (< 2 m) water within the lagoon of Rarotonga (Cook Islands; 21°13’38”159°46’34”W; May 9^th^-15^th^ 2023). Colonies were selected at random across four sites (Āroa, Avarua, Betela, and Nikao; n = 12 or 13 per site). From each colony, five samples (each comprising 2-3 individual monticules) were collected using a 3 mm diameter hole punch and hammer, at regular intervals from colony apex to base along the southern face (Figure 1). This sampling region was expected to span a high to low light gradient created by colony self-shading, reflecting the primarily northern sunlight direction at Rarotonga’s South Pacific location. Orientation was determined using a compass, levelled on the benthos with a handheld spirit level prior to sampling alongside a coded target tile for digital 3D reconstruction (see Supplementary Methods). Non-toxic epoxy putty was used to seal sampling sites and indicate intra-colony sample position (Knead It Aqua, Selleys). Samples were immediately rinsed in 0.22 µm filtered seawater and stored in DMSO buffer [57] for subsequent DNA extraction. Colony depth was determined as distance from the water’s surface to the first sample position (colony apex) at low tide.

**Figure 1.**
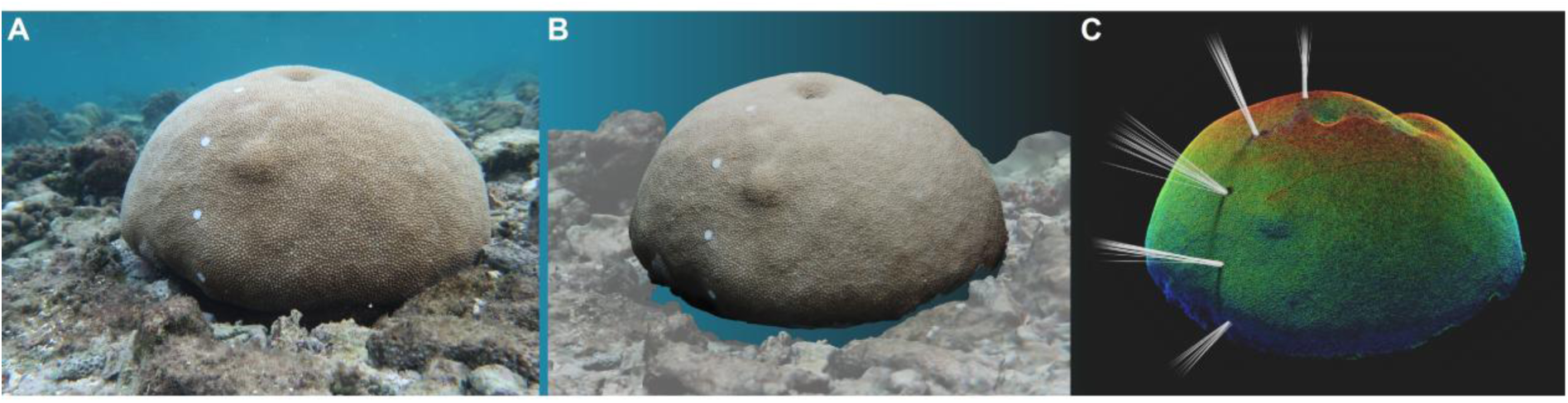
**A)** Image of a *Hydnophora microconos* colony, with white epoxy circles indicating sample positions (1-5; apex to base). Colony height range (distance between apex-most and base-most points) was 21.55 cm. **B)** Digital three-dimensional reconstruction of sampled colony, isolated from surrounding reef geometry. **C)** Digital three-dimensional reconstruction of isolated colony coloured by per-vertex ambient occlusion. The z normal direction (upwardness) is displayed as white lines stemming from vertex clusters at each sample position.

### Photogrammetry and 3D mesh processing

Between 75 and 150 images with approximately 70% overlap were captured of each colony and its proximate substrate for 3D photogrammetric reconstruction (Canon G7xii in a Nauticam housing). Three-dimensional models were reconstructed via Agisoft Metashape (Professional Edition, version 2.0.2) using script-based batch processing. Vertex confidence was visually reviewed (see Supplementary Materials), with particular focus on sample positions as a qualitative indicator of reconstruction accuracy. A workflow developed for effective post-build processing using the software Meshlab (v2023.12) and R (version 4.3.1 [58]) was subsequently applied to all colony reconstructions. Full model processing workflow is provided as Supplementary Methods.

### Morphological quantification of colonies

Three key aspects of geometric structure were quantified, using six paired morphology metrics: 1) volume compactness (sphericity and convexity); 2) top-heaviness (second moments of area and volume); and 3) surface complexity (packing and fractal dimension) (see Supplementary Methods) following methods developed by Zawada et al. [50]. All calculations were performed using the R package Habtools [59], designed for extracting structural information from 3D objects. The magnitude variables surface area, volume, and height range were additionally defined from isolated colony meshes (Rvcg [60]). Morphology metrics, and their calculation, are detailed at length elsewhere [50,51,59], but are described briefly below to support interpretation.

Convexity quantifies volume compactness by comparing an object’s volume to that of its convex hull, whilst sphericity evaluates how closely an object’s geometry approaches a sphere. Second moments of area and volume assess vertical distribution, indicating top-heaviness. Surface complexity was characterised through packing, which compares an object’s surface area to that of its convex hull, alongside fractal dimension, which measures the space-filling capacity of an object.

### Sample-scale spatial analysis

Vertex clusters were generated with a 3 (± 0.3) mm radius at each sample location (Meshlab). Sample location in three-dimensional space was determined by deriving average x, y, and z coordinates from corresponding vertex clusters, with sample depth indicated by position along the z axis. Average z normal direction (hereafter ‘upwardness’) was also derived for vertex clusters to inform sample orientation (Figure 1).

Additionally, ambient occlusion models were generated for each processed 3D scene (Figure 1). This was achieved using Meshlab’s *Ambient Occlusion* function, a per-vertex calculation that evaluates the extent to which model geometry is exposed to ambient light. Light input was simulated along the z axis (x, y, z lighting direction: 0,0,1). Directional lighting bias was set at 0.8, approximating minimal light scattering expected within the top 2 metres of the water column [61,31,62,63]. Default inputs were retained for all other settings. Per-vertex ambient occlusion values were mapped to sample vertices by aligning corresponding vertex coordinates, allowing the characterisation of ambient occlusion at sample-scale.

### Symbiodiniaceae identification & phylogenetic analysis

DNA was extracted from all samples following a modified CTAB-chloroform protocol, and the internal transcribed spacer 2 (ITS2) ribosomal region amplified through PCR [45]. Amplicons were sequenced on an Illumina MiSeq platform (Auckland Genomics), with raw sequence data submitted to SymPortal [64] and SymPortal outputs used for symbiont characterisation. To validate putative *Cladocopium* taxa detected by SymPortal, congruence of the plastid minicircle psbA non-coding region (psbA^ncr^) with ITS2 type profiles was tested for across a subset of samples. See Supplementary Material for full molecular biology methods.

ITS2 sequence similarity was assessed across samples for all major genera identified. A k-mer-based similarity matrix was generated using ‘*kdistance’* (k = 6) (kmer [65]), from which unifrac distances with an alpha of 0.5 were calculated (*unifrac*, GUniFrac [66]), with output visualised alongside per-sample sequence and ITS2 type profile composition [17,67].

psbA^ncr^ sequences were manually trimmed following visual inspection and aligned using AlignSeqs (DECIPHER [68]) to generate consensus sequences; sequences of poor alignment were first removed from downstream analysis. Reference sequences from recent *Cladocopium* C3-radiation species descriptions [20] were aligned with psbA^ncr^ sequences (*AlignSeqs*, DECIPHER). The position of the study *Cladocopium* sequences relative to reference sequences was investigated using four different tree-building methods: 1) maximum parsimony, 2) maximum parsimony with insertions and deletions coded as a 5^th^ character state, 3) maximum likelihood, and 4) alignment-free k-mer with jack-knifing (see Supplementary Methods); relationships were inferred from method consensus.

Linear deconvolution (non-negative least squares) using psbA^ncr^-validated reference ITS2 profiles was used to detect samples where SymPortal had likely merged *Cladocopium* sequences from co-occurring taxa into a single meta-profile, otherwise masking *Cladocopium* co-occurrence (see Supplementary Methods).

### Statistical analysis

All analyses were conducted in R version 4.3.1 [58]. Code is accessible at GitHub and Zenodo (link will be updated upon acceptance to journal).

Relative ITS2 sequence abundances were summed by majority genera. Each colony and sample were individually categorised as dominated by a single symbiont genus (≥ 98% of sequences belonging to a single genus) or hosting a multi-symbiont assemblage. Non-parametric one-way analysis of variance (Kruskal-Wallis) and *post hoc* Dunn Tests assessed site-level differences in symbiont genera and colony depth.

Principal Components Analysis (PCA; *rda*, vegan [69]) evaluated colony morphology based on six morphometrics (see above) together with colony height range. Symbiont composition was tested against the ordination of colony morphology within multivariate space using permutational analysis of variance both overall (PERMANOVA) (*adonis2*, vegan) and between composition types (*pairwise.adonis*, pairwiseAdonis [70]). Homogeneity of multivariate dispersion of symbiont compositional groups was tested using the *betadisper* function (vegan). Paired t-tests were also used to compare significant morphometrics individually against symbiont composition.

Stepwise detection analysis was utilised to determine the likelihood of detecting multi-symbiont assemblages based on incremental sample inclusion from colony apex to base, as well as the sampling approaches of random sample selection, colony apex plus a randomly selected sample, and colony apex plus base [71,45,72]. Across 100 iterations, per-colony genera prevalence was determined for each sampling approach using random resampling with replacement of ITS2 sequences (n = 1500), weighted by relative abundance. The detection of multiple symbiont taxa was then denoted according to a 98% single genus threshold (see above). A binomial generalised linear mixed model (GLMM) was subsequently run to evaluate community detection rate against the interactive fixed effects of sampling approach and the number of colonies sampled (n = 1-49), with iteration held as a random effect (glmmTMB [73]). Estimated marginal means (emmeans [74]) subsequently estimated the implied probability of community detection for each sampling approach, across each number of included colonies. Pairwise comparisons were generated for community detection across all sampling approaches, with a Holm adjustment for multiple comparisons (*contrast,* emmeans).

Zero-inflated, negative binomial GLMMs were used to explore per-sample genus abundance against each of the fixed effects of sample depth, upwardness, and ambient occlusion, with colony-specific zero-inflation terms and colony identity as a random intercept (glmmTMB).

## Results

### Hydnophora microconos hosts diverse Symbiodiniaceae

Sequencing of the ITS2 gene region yielded approximately 4.5 million Symbiodiniaceae sequences across all samples. The per-sample average sequencing depth was 18,623 (± 5978 SD), following removal of one sample with low read count (< 1500). Over 99% of identified sequences belonged to the genera *Cladocopium* (67.59%) and *Durusdinium* (32.20%). Approximately 0.21% of sequences, attributable to other genera, were excluded from subsequent analysis.

SymPortal analysis identified 23 ITS2 type profiles, largely characterised by the majority defining intragenomic variants (DIVs) ‘C3/C3f’, ‘C3/C3k’, or ‘D1’. Among ITS2 profiles identified, C3/C3f-C3b-C3.10-C3fl-C3gj-C3ae-C21 occurred most frequently (26% of total profiles), whilst C3/C3k-C50a-C3.10-C3gj-C3b, D1-D4-D6-D1b-D1ab-D6m, and D1-D4-D6-D1b-D1d-D6c-D10d each represented an approximate further 15% of total profiles. Approximately 23% of sequences did not contribute to ITS2 profiles.

A hierarchical clustering of generalised UniFrac distances across *Cladocopium* sequences revealed clear division between C3/C3k and C3/C3f ITS2 type profiles (Figure S1). Sequencing of the psbA^ncr^ for representative sets of C3/C3k and C3/C3f samples demonstrated distinct monophyletic lineages that were completely congruent with ITS2 profile data. All tree-building methods positioned these lineages as most closely related to *Cladocopium sodalum* (Figure S2) [20]. Hereafter, these lineages are referred to as ‘C3f’ and ‘C3k’. Equivalent divisions were not present in sequences identified within the genus *Durusdinium* (Figure S1), and we hereafter assume there to be a single *Durusdinium* taxon present (‘*Durusdinium’*).

Linear deconvolution analysis revealed five samples with intermediate contributions from C3f and C3k reference profiles (0.4-0.6), indicating that these samples were poorly explained by either reference profile alone and instead showed shared C3f-C3k characteristics (see Supplementary Material).

### Cladocopium and Durusdinium co-occurred across sites

*Cladocopium* and *Durusdinium* were similar in abundance across the sites Āroa, Betela, and Nikao, where *Durusdinium* comprised 32-53% of sequences (Table 1). The site Avarua was distinguished by substantially increased *Cladocopium* prevalence (Kruskal-Wallis test: χ² = 45.034, p < 0.001 | *post hoc* Dunn test: p < 0.01), with around 5% of sequences belonging to *Durusdinium* (Table 1). C3k ITS2 profiles were not observed at Avarua.

**Table 1.** Sampling summary, library statistics, and symbiont composition of *Hydnophora microconos* across sites. Labels ^a^ and ^b^ denote *post hoc* Dunn Test grouping.

|  | Āroa | Avarua | Betela | Nikao |
| --- | --- | --- | --- | --- |
| N colonies | 13 | 12 | 13 | 12 |
| N samples | 61 | 58 | 62 | 60 |
| Average per-colony sequencing depth (± SD) | 80,332 (± 29,462) | 94,461(± 15,825) | 87,000 (± 16,174) | 98,276 (± 17,002) |
| Lat \| Lon | -21.25570 \| -159.8195 | -21.20394 \| -159.7696 | -21.24610 \| -159.8262 | -21.20763 \| -159.8243 |
| *Cladocopium* / *Durusdinium* ratio | 0.679/0.321<sup>b</sup> | 0.957/0.043<sup>a</sup> | 0.475/0.525<sup>b</sup> | 0.602/0.398<sup>b</sup> |
| Number of *Cladocopium*-dominated colonies | 2 | 7 | 0 | 2 |
| Number of *Durusdinium*-dominated colonies | 2 | 0 | 4 | 3 |
| Number of colonies with communities | 9 | 5 | 9 | 7 |
| Average colony depth (cm) (± SD) | 76 (± 14)<sup>b</sup> | 27 (± 4)<sup>a</sup> | 62 (± 20)<sup>b</sup> | 66 (± 26)<sup>b</sup> |

Average colony depth was significantly different across sites (Kruskal-Wallis test: χ² = 23.928, p < 0.001). Average colony depth at Avarua (27 cm) was approximately 2.5 times shallower than at Betela and Nikao and 2.9 times shallower than at Āroa (Table 1).

### Inter-colony symbiont composition was heterogeneous

Dominance by *Cladocopium* or *Durusdinium* was identified in 40% of colonies (n = 11 and n = 9, respectively), whereas 60% of colonies (n = 30) hosted a multi-symbiont assemblage (Figure 2, Figure 3). Of colonies that were *Cladocopium* dominated, 10 colonies were dominated by C3f. Multi-symbiont assemblages were characterised according to three co-occurrence types: C3k and *Durusdinium* (C3k|D), C3f and *Durusdinium* (C3f|D), or C3k, C3f, and *Durusdinium* (C3k|C3f|D). Notably, the most prevalent symbiont composition was C3k|D, occurring in 28% of all colonies, although dominance by C3k alone was rare (n = 1). Given this low representation of C3k dominance, it was removed from morphometric analyses (see below). C3f|D was identified in 22% of colonies, whilst 10% of colonies were associated with all three major taxa C3k|C3f|D. In the absence of *Durusdinium,* C3k and C3f co-occurrence was not observed.

**Figure 2.**
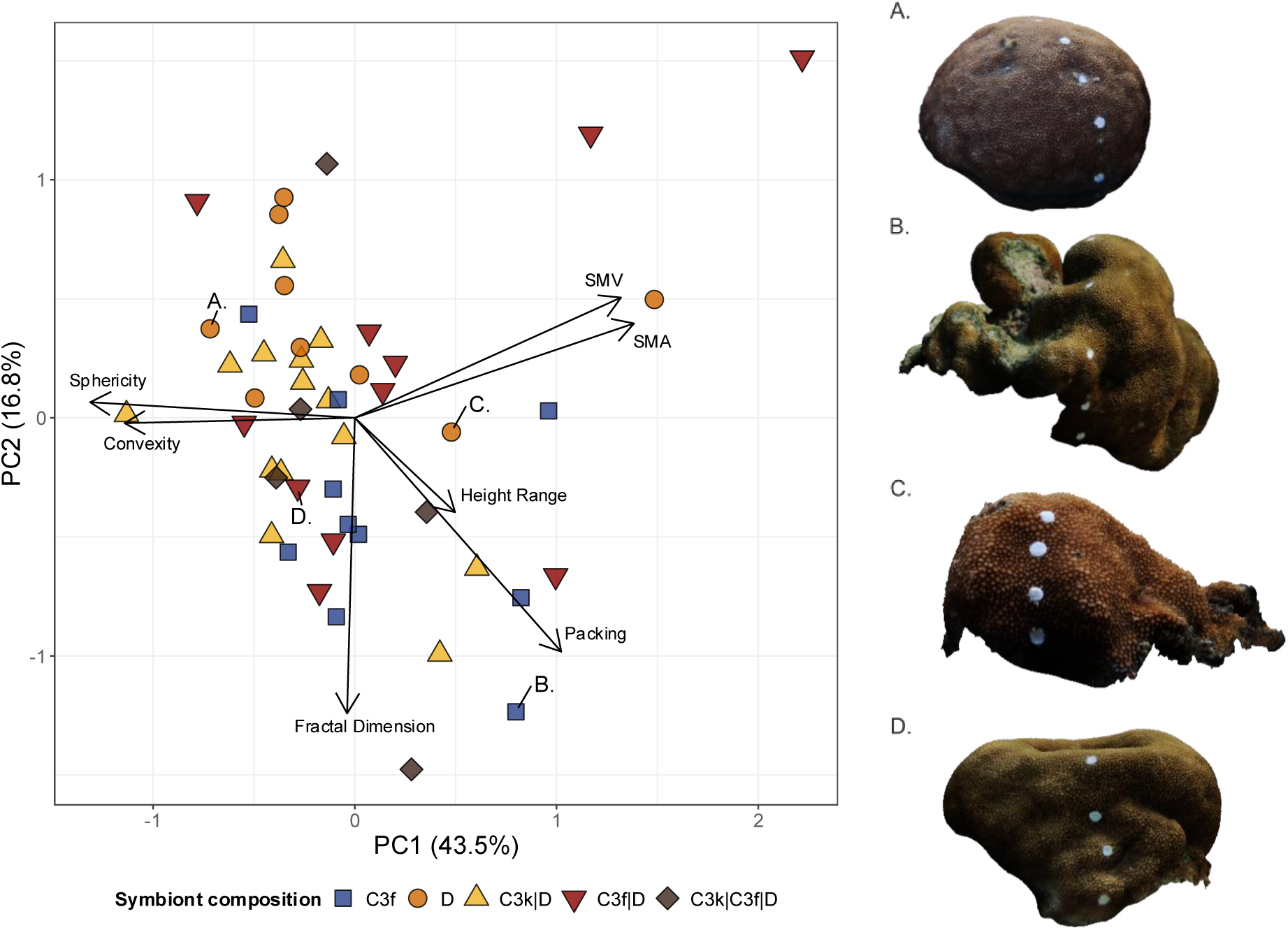
Principal Components Analysis of colony morphometrics. Each point represents one *Hydnophora microconos* colony. Points are coloured and shaped according to intra-colony symbiont composition. Labelled points (ABCD) indicate the morphometric distribution of right-hand panel colonies.

**Figure 3.**
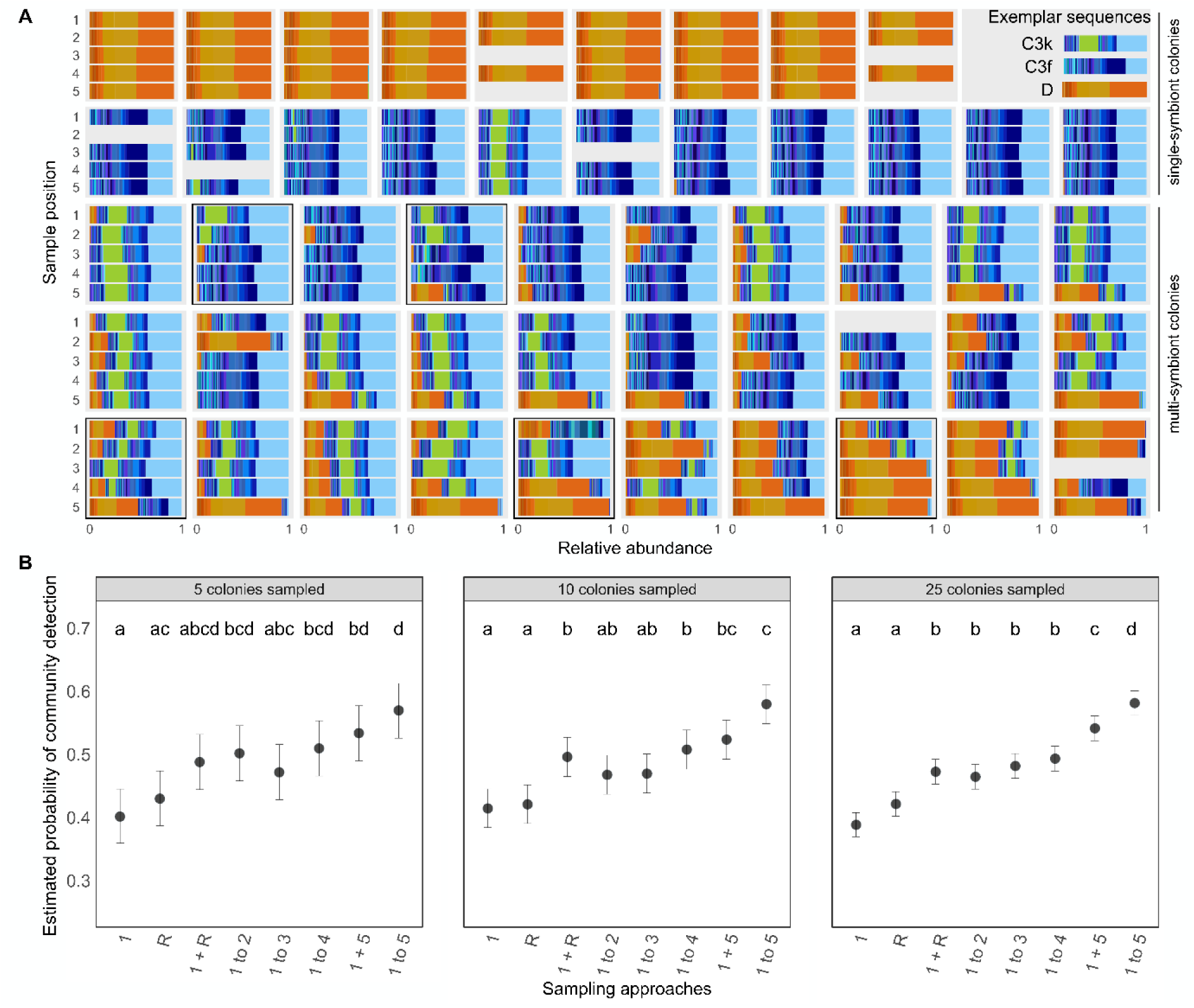
**A)** 50 colonies highlighting the sample positions, genetic identity, and community structure of each colony from apex (position 1) to base. Exemplar sequences indicate C3k, C3f, and *Durusdinium*(D) ITS2 sequence composition. Black boxes surround colonies with C3k|C3f|D co-occurrence. **B)** Estimated probability of colony-level *Cladocopium-Durusdinium* co-occurrence detected across sampling approaches (± asymptotic lower and upper confidence limits), for 5, 10, and 25 colony sampling scenarios. Letters denote pairwise grouping.

### Coral colonies are morphologically diverse

Colony size ranged along a morphological continuum. Surface area extended from 590 cm^2^ to 7087 cm^2^ (mean: 1748 ± 166 cm^2^ SE), reflecting an approximate 12-fold difference between the least and greatest host tissue area, whilst volume spanned 576 cm^3^ to 36,322 cm^3^ (mean: 6913 ± 933 cm^3^ SE). Height range, represented by distance along the z axis from the minimum to maximum vertex, varied from 10.4 cm to 43.5 cm (mean: 21 ± 0.9 cm SE).

PCA revealed a broad distribution of colonies within multivariate morphometric space. The first two Principal Components (PC) captured 60% of the observed variation in colony geometry (Figure 2). PC1 (43.5%) reflected differentiation in volumetric efficiency from compact, convex colonies to convoluted, top-heavy forms, influenced by strong positive loadings from the morphology metrics packing and second moments of area and volume, together with negative contributions from sphericity and convexity; PC2 (16.8%) was largely driven by the surface complexity variables fractal dimension and packing, with slight influence from height range (Figure 2).

### Symbiont community composition is related to colony morphometrics

Sixteen percent of the variance in colony distribution within multivariate morphometric space was explained by symbiont composition (PERMANOVA, F_4,44_ = 2.153, p < 0.05), but not site (PERMANOVA, F_3,45_ = 0.971, p = 0.449; Figure S3), with variance similar across groups (betadisper, F_4,44_ = 0.844, p = 0.51, permutations = 999). Colonies dominated by *Durusdinium* were morphologically distinct from those dominated by C3f (pairwise PERMANOVA, adjusted p < 0.05). This distinction occurred primarily along the PC2 axis, where 70% of C3f-dominated colonies were positioned within the negative PC2 space, categorised by high fractal dimension and packing, whereas colonies dominated by *Durusdinium* largely occupied positive PC2 values (see Figure 2). Concordantly, mean packing and fractal dimension values were significantly greater in C3f-dominated colonies than in those dominated by *Durusdinium* (Welch’s t-tests: both p < 0.05). Height range trended toward significance, with *Durusdinium-*dominated colonies clustering toward lower height range values (Welch’s t-test: p = 0.057). Colonies hosting the co-occurring symbionts C3k|D, C3f|D, and C3k|C3f|D did not exhibit distinct clustering and broadly intersected the distribution of single taxon colonies (pairwise PERMANOVA, adjusted p < 0.05) (see Figure 2).

### Intra-colony sampling enhances co-occurrence detection

Multi-symbiont assemblages were observed across all sample positions, with *Cladocopium-Durusdinium* co-occurrence identified in 42% of samples overall. The highest proportion of samples with *Cladocopium-Durusdinium* co-occurrence occurred at the colony base (sample position 5) (Table S5). Co-occurrence of C3k|D and C3f|D were observed in similar frequency across all samples, whilst C3k|C3f|D was not detected at the sample level; full library statistics across sample positions are provided in Table S5.

Stepwise detection analysis was carried out to evaluate the influence of variable sample inclusion on *Cladocopium-Durusdinium* community detection. A single colony where sample position 1 was not available was excluded from this analysis. Community detection was lowest when sampling the colony apex in isolation, or when sampling at random (R), and greatest when intra-colony sampling along an apex-to-base gradient (Figure 3). This overall trend in community detection was consistent across an increasing quantity of included sampled colonies, yet sampling approaches became increasingly differentiated as colony number increased (Figure 3). For example, when sampling fewer colonies (n = 5 or n = 10), targeting colony apex and base alone (positions 1 + 5) was equivalent to sampling the complete vertical gradient (positions 1 to 5), whereas when sampling 25 colonies, sampling all five positions emerged as the strongest technique (Figure 3).

### Intra-colony symbiont distribution is related to spatial position

Variance in upwardness increased from colony apex to base (Figure 4), capturing the complexity of colony structures. Upwardness reflected near vertical orientation at the colony apex (median: 0.95), whilst 65% of samples were oriented negatively along the z axis at the colony base (median: -0.271) (Figure 4). Likewise, ambient occlusion differed substantially across the colony surface (Kruskal-Wallis test: χ² = 95.422, p < 0.001) and was lowest at the colony base (mean: 0.075 ± 0.022 SE), increasing exponentially (Figure S4) with an approximate 10-fold increase at the colony apex (mean: 0.738 ± 0.01 SE) (Figure 4).

**Figure 4.**
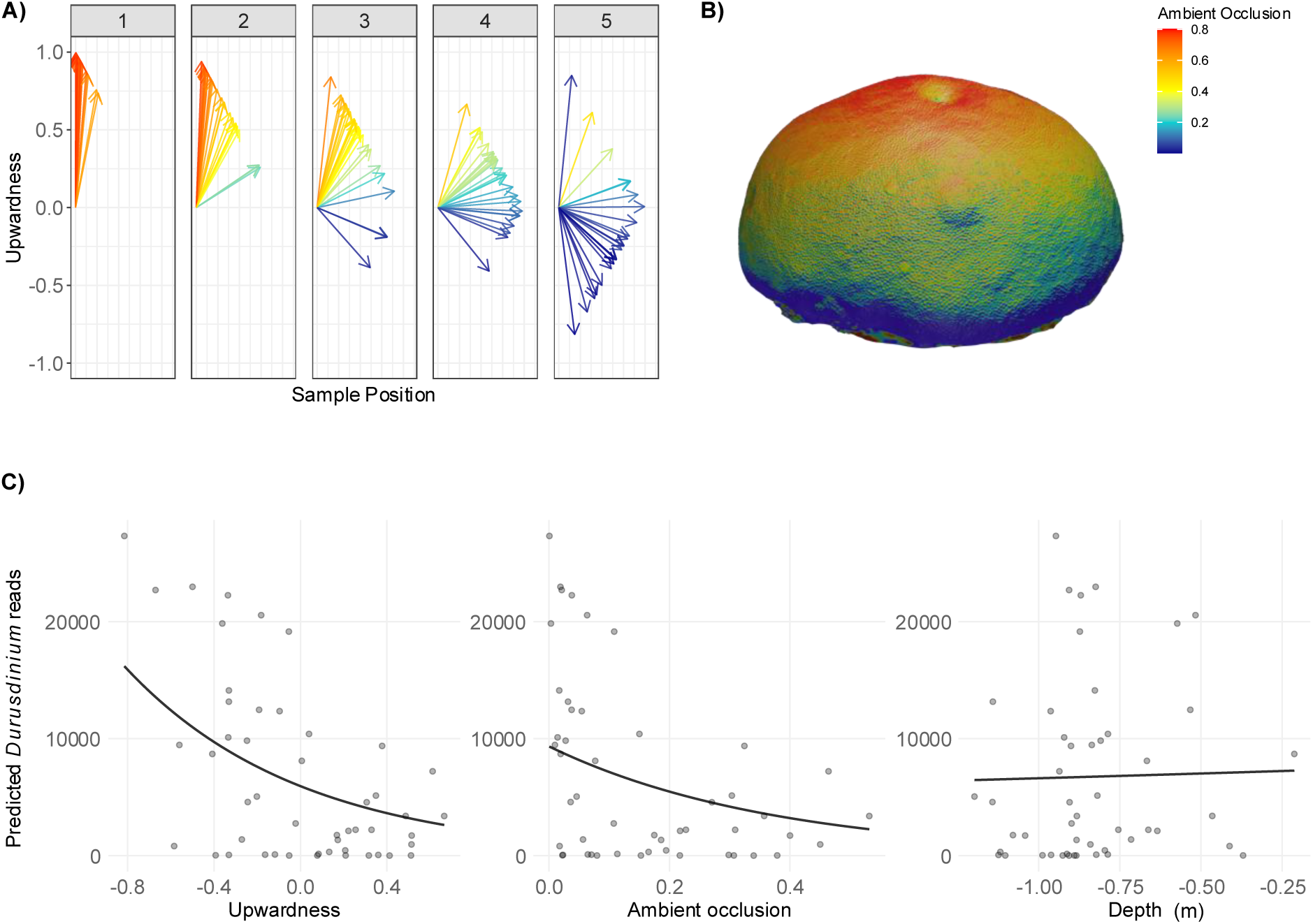
**A)** Per-sample upwardness across intra-colony sample positions (apex-base: 1-5) for colonies with *Cladocopium-Durusdinium* assemblages, excluding the site Avarua (n colonies = 25, n samples = 125). Each arrow represents a sample, coloured by its corresponding ambient occlusion value. **B)** Example colony displaying ambient occlusion as a colour gradient. **C)** Zero-inflated negative binomial generalised linear mixed models for the variables ‘upwardness’, ‘ambient occlusion’, and ‘depth’, predicting *Durusdinium* read count across the colony base region (sample positions 4 and 5). Models included colony-specific zero-inflation terms and colony as a random intercept. Points represent individual samples; the black line represents the model predicted reads. Models excluded the site Avarua and a single sample outlier.

The parameters ‘upwardness’, ‘ambient occlusion’, and ‘sample depth’ were individually modelled against per-sample *Durusdinium* abundance using zero-inflated negative binomial GLMMs. Full models incorporated colonies where multi-symbiont assemblages were present, excluding the distinctly shallow Avarua site that exhibited near complete *Cladocopium* dominance (n = 25). Full models demonstrated a negative relationship between *Durusdinium* abundance and per-sample upwardness (slope = -0.6736, p = 0.02) as well as ambient occlusion (slope = -1.1532, p = 0.02), whilst sample depth did not predict *Durusdinium* abundance (p > 0.05) (Figure S5). Reduced models that excluded the top-most colony region (sample positions 1-3) alongside an outlier sample increased the strength of this relationship. *Durusdinium* abundance increased from sample position 4 to 5 in relation to decreasing upwardness and ambient occlusion (slope = -1.2312 and -2.6747, respectively, p < 0.05), with sample depth remaining an insignificant predictor (Figure 4). Where the outlier sample (upward-oriented, highly occluded base sample, Figure 4, Figure S4) was included, ambient occlusion predicted *Durusdinium* abundance, whereas upwardness failed to capture this trend (Figure S5).

## Discussion

### Hydnophora microconos is a model for investigating intra-colony Symbiodiniaceae co-occurrence

*Hydnophora microconos* has been proposed as a model species for studying multi-symbiont associations in the Pacific [45]. The comprehensive sampling assessment presented here characterised frequent *Cladocopium*-*Durusdinium* co-occurrence across colonies and sites, supporting the suitability of this species as a model system. High spatial resolution afforded by the present sampling design revealed an association with C3k not captured previously at the study location [45], with C3K|*Durusdinium* representing the most common form of co-occurrence. To our knowledge, this is the first study to apply the ITS2-psbA^ncr^ congruence approach to intra-colony samples, revealing the co-occurrence of *Cladocopium* lineages (C3k and C3f) within colonies. Examples of *Cladocopium-Cladocopium* interactions in colonies are exceptionally rare [45]. Further work is required to resolve C3k-C3f genetic distances (i.e., genotypes vs. species) and functional biology to understand how closely related lineages coexist within coral colonies. Overall, the prevalence of structured intra- and inter-genus symbiont communities across variable, geometrically complex colonies broadens the utility of this model, positioning *H. microconos* as a strong system for evaluating the influence of colony architecture on the ecological processes that structure symbiont communities.

### Colony geometric complexity was associated with Cladocopium-Durusdinium co-occurrence

We applied methods, previously developed by Zawada et al. [50,51] for quantifying coral morphology across growth forms amongst the order Scleractinia, to evaluate morphological variation within a single coral taxon. This represents a fine-scale application of high-resolution photogrammetry, where comparatively subtle geometric variation between colonies may have been expected to constrain morphological resolution. *H. microconos* is traditionally described as massive in growth form [54,55], although the species is also known to be encrusting [56] (Figure S6). Relative to other broad-scale growth forms, massive and sub-massive colonies are expected to exhibit high volume compactness, constraining surface complexity and top-heaviness [50]. Correspondingly, massive colonies show low morphological complexity and reduced inter- and intra-specific morphological variation [52]. Within these outlined geometric bounds, however, *H. microconos* colony structure showed substantial geometric spread across multivariate morphometric space. Moreover, there was sufficient structural differentiation to reveal an association between symbiont composition and geometry, whereby distinct geometries were resolved for colonies dominated by C3f or *Durusdinium*. Application of this approach to other morphologies, such as encrusting-branching, in addition to the encrusting-massive morphology studied here, is required to understand the broader relevance of geometry.

Morphometric distinction between single taxon colonies primarily reflected a gradient of high to low surface convolution. Although site-specific environmental variation can influence coral form [75–79], and C3f dominance was heightened at the distinctly shallow Avarua site, site was not a significant predictor of morphology, nor depth a significant predictor of symbiont prevalence. Reductions in the geometry metrics packing and fractal dimension signify decreased surface convolution in *Durusdinium-*dominated colonies, additional to a trend toward reduced height range. Several studies point toward trade-offs in coral form and function through association with distinct Symbiodiniaceae genera (but see [80,81]). In particular, when in symbiosis with *Durusdinium* (versus *Cladocopium*) corals appear to experience reduced carbon translocation from the symbiont [9], reduced calcification [82], and/or a reduced growth rate [83–85]. An influence of symbiont identity on host growth rate would likely indicate a pathway for symbiont-mediated impact on colony geometry, driving intra-specific morphological difference in corals with differing symbionts.

Where colonies showed *Cladocopium*-*Durusdinium* co-occurrence, distinct geometries were not associated with symbiont composition, with colonies reflecting the entire range of geometric complexity. The modular nature of coral organisms dictates that colony growth requires individual polyp replication, where polyp addition rate is linked to the rate of skeletal deposition [33]. Polyp replication is variable across the colony surface [33], suggesting an absence of spatial uniformity in colony growth. Consistent with this observation, skeletal height at the polyp level is reduced under asymbiotic conditions [86], further indicating that skeletal modification is influenced by polyp-specific physiology. As such, host growth may be modulated, at least in part, at the level of individual units. If this notion is indeed correct, any influence of symbiont identity on geometric complexity would likely be enhanced where multiple symbionts are hosted, with differing symbiont contribution to host growth supporting variation in host form.

### Sampling design is important when quantifying multi-symbiont prevalence

It has been suggested by recent meta-analyses of *Cladocopium* and *Durusdinium* co-occurrence that the lack of systematic intra-colony sampling has likely resulted in a field-wide under-estimation of the prevalence of symbiont co-occurrence [41]. In our current study, stepwise detection analysis revealed the utility of spatially informed, intra-colony sampling for detecting multi-symbiont assemblages within colonies. Community detection was particularly strong where an entire apex to base gradient along the colony surface was considered, however optimality of this approach was exemplified upwards of around 25 individual colonies. This finding reflects site-level differences but also uneven distribution of *Cladocopium* and *Durusdinium* across *H. microconos* colonies, and is indicative of niche differentiation as if both genera were uniformly distributed, random sampling, or sampling the apex alone, would have been sufficient to accurately quantify symbiont community composition. Consequently, efficiently detecting patterns at relevant scales likely requires sampling strategies able to capture underlying spatial structure in symbiont distributions. Field studies employ a range of sampling approaches, ranging from position-standardised single samples when maximising for the number of colonies sampled [87], to a fixed plus random sample approach [72] or targeting intra-colony extremities [45,88]. Incorporating sensitivity or power analyses into future work would aid in optimising sampling effort for enriched realisation of intra-colony symbiont diversity.

### Morphologically driven niches predict Durusdinium prevalence

Host microanatomical features modify the local tissue environment at scales relevant to individual symbiont cells due to boundary-layer effects [89,90], with higher-order modulation of processes such as current and light exposure by macro-scale colony structures [89,91,92]. Whilst variable host morphology represents an overarching source of heterogeneity, a geometric aspect that is conserved across scales is ‘monticule (i.e., the sample) inclination’, oriented normal to the colony surface. Photogrammetric models preserved true-to-life orientation and scale, enabling comparable evaluation of three-dimensional positioning across samples. Strong intra-colony light gradients are documented, where scalar irradiance at the polyp surface substantially decreases from the exposed (apical) to shaded (base) regions [31]. Specialisation to microenvironmental conditions can occur at the polyp level [93]. Furthermore, intra-colony stratification of symbiont physiology has been observed, where along an intra-colony light gradient, low light exposed symbionts have comparatively increased photosynthetic efficiency and starch storage [94]. Such microenvironmental variation and resulting responses of symbiont functional biology present a strong case for sufficient heterogeneity over which niche partitioning can occur. We identified such partitioning whereby *Durusdinium* increased in prevalence at the often underhanging base region of colonies, where modelled ambient occlusion was greatest. However, for corals exhibiting complex geometries, upwardness was not always a strong proxy of occlusion. Notably, ambient occlusion captured a trend in *Durusdinium* reduction where upwardness was insufficient, owing to a rare, upward-oriented, highly exposed base sample that was dominated by *Durusdinium* (Figure S4), highlighting the benefit of accounting for multiple environmental contexts alongside patterns in symbiont distribution.

Members of the genus *Durusdinium* are frequently referred to as extremophiles or opportunists, known for increased prevalence in atypical reef environmental conditions including elevated thermal environments [95], but also cooler environments [96], areas of high turbidity [16], and after bleaching [97]. The increase in *Durusdinium* in light-occluded regions proximate to the benthos may suggest microenvironmental suitability related to its opportunistic attributes. Although it was not the case that *Durusdinium* was excluded from highly exposed, apex regions, prevalence increased where occlusion was strongest, indicating that the apex and side of the colony may represent a region of increased competition, where *Durusdinium* prevalence becomes more challenging to predict. The degree to which these upper colony regions represent stable states of *Cladocopium-Durusdinium* competition is unknown, yet the orientational, and thus exposure, difference at the colony base is indicative of diverging microenvironmental conditions where host-symbiont optimality is better favoured through a *Durusdinium* partnership.

## Conclusion and future directions

Symbiont genetic and functional diversity has been implicated as a major determinant of coral fitness, yet the mechanisms facilitating distinct symbiont co-occurrence are poorly resolved. In particular, the contribution of coral colony architecture towards symbiont co-occurrence remained unquantified. Here we demonstrate that distinct Symbiodiniaceae exhibit spatially structured distributions within geometrically complex coral colonies. Given that colony morphology, itself broad and continuous across the population, is clearly linked to symbiont community composition, random sampling of only a few colonies underestimates this structure, and only a pooled, intra-colony sampling approach can adequately resolve how morphospace shapes symbiont niche differentiation. Future work characterising temporal stability of intra-colony distributional patterns and evaluating the extent to which symbiont physiology may drive morphological complexity of host coral through controlled experimentation, will build insight into the role of multi-symbiont assemblages in host performance.

## Acknowledgements

We express our gratitude to Guy and Glenda Nitschke whose assistance with field collections made this research feasible.

