## Supplementary_V1 for "Colony geometric complexity predicts the co-occurrence of coral photosymbiont genera"

### Supplementary methods and materials

#### *Photogrammetry and 3D mesh processing*

Each model was first size calibrated and levelled in the xy-plane in Agisoft Metashape (Professional Edition, version 2.0.2) via detection of coded targets, positioned at known distances on an in-scene marker tile. Confidence was displayed across the mesh as a colour map and visually reviewed, ensuring high confidence at locations of sample collection. Meshes exported with associated per-vertex confidence values are available at Zenodo (link to be updated upon acceptance to journal). The software Meshlab (v2023.12) was then used to transform the x, y, and z axis coordinates to achieve true-to-life orientation of the model. Following initial alignment of the positive y axis to magnetic north against an in-scene reference compass, the point of origin was set where x and y axes intersected the centre of the first sample position (i.e., the colony apex), and adjusted to colony-specific depth along the z axis. All non-colony components were then removed to generate an isolated colony model. An additional closed "watertight" version of each isolated colony model was generated using Meshlab's *Close Holes* function to satisfy requirements for geometric calculations. All subsequent processing was carried out in the statistical environment R (4.3.1 [1]) using the package Rvcg [2]. To support cross-colony morphological compatibility, meshes were uniformly remeshed with a minimum resolution of 1 mm. All final meshes were visually inspected for morphological integrity.

#### *Quantifying colony morphology*

For the evaluation of colony morphology, morphometric values were first transformed following methods outlined in Zawada et al. [3] including logit transformations for sphericity and convexity values and log10 transformations for packing and second moments of area and volume. All metrics were subsequently standardised to a zero mean and unit variance (*decostand*, vegan [4]).

Surface area is central to the metrics sphericity and packing. *H. microconos* colonies were highly varied in their attachment style, with some colonies having formed on the side of existing rock/reef structures (i.e., encrusting) whilst others were situated on the benthos. Consequently, following isolation from the substrate, the extent of the "open" portion across colony meshes was inconsistent. This variability introduced an attachment bias with potential to influence cross-colony compatibility. To promote consistency, measurements of sphericity and packing were obtained from closed "watertight" versions of each colony mesh.

#### DNA extraction and sequencing

#### ITS2 sequencing and Symbiodiniaceae identification

The internal transcribed spacer 2 (ITS2) ribosomal gene region was amplified through PCR using the Symbiodiniaceae-specific primers SYM_VAR_5.8S2 (TCGTCGGCAGCGTCAGATGTGTATAAGAGACAGGAATTGCAGAACTCCGTGAACC) and SYM_VAR_REV (GTCTCGTGGGCTCGGAGATGTGTATAAGAGACAGCGGGTTCWCTTGTYTGACTTCATGC) (including Illumina adaptor sequences). Optimized PCR conditions included initial denaturation for 2 minutes at 95°C, followed by 35 cycles of 95°C for 45 seconds, 68°C for 45 seconds, and 72°C for 30 seconds, followed by a final extension at 72°C for 5 minutes. Following the manufacturer's protocol, normalised amplicons underwent a second indexing PCR using Nextera XT Index primers. Samples were purified twice under a 1:1 ratio of AmpureXP beads following equimolar pooling. The final library was sequenced on an Illumina MiSeq platform using a 2 x 300 bp V3 kit with the addition of approximately 17% phiX (Auckland Genomics).

Raw sequencing data were submitted to SymPortal [5], an analytical framework that recognises within-sample intragenomic sequences, referred to as 'defining intragenomic variants' (DIVs), and characterises taxonomically distinct Symbiodiniaceae through patterns of DIV combination. SymPortal identifies distinct Symbiodiniaceae through ITS2 type profiles, denoted by their respective collections of DIVS in order of descending abundance [5]. This approach resolves complications in interpreting Symbiodiniaceae diversity otherwise presented by the multicopy nature of the ITS2 marker. SymPortal DIV and ITS2 profile output were used for symbiont characterisation.

**Sequencing of the psbA$^{ncr}$**

To amplify the psbA non-coding region (psbA$^{ncr}$) across a subset of *Cladocopium* dominated samples, DNA was extracted from an aliquot of homogenised tissue following a modified CTAB-chloroform protocol. PCR then used the primer pair 7.4-For (5` - GCATGAAAGAAATGCACACAACTTCCC – 3`) 7.8-Rev (5` - GGTTCTCTTATTCCATCAATATCTACTG – 3`) that preferentially amplify psbA$^{ncr}$ haplotypes from the *Cladocopium* lineage [6]. Optimised PCR conditions involved an initial denaturation step at 95°C for 2 minutes, followed by 35 cycles of 94°C for 10 seconds, 58°C for 30 seconds, and 72°C for 2 minutes, followed by a final extension of 72°C for 10 minutes. PCR amplicons were sequenced in both forward and reverse directions (Sanger Sequencing, Macrogen Inc. South Korea).

Reference sequences (GenBank OQ359894–OQ359908, OQ359909–OQ359921, and OQ359922–OQ359927) from recent *Cladocopium* C3-radiation species descriptions [7] were aligned with psbA$^{ncr}$ sequences (*AlignSeqs*, DECIPHER [8]), with aligned sequences end-trimmed to remove overhangs. The position of the study *Cladocopium* sequences relative to reference sequences was then investigated using four different tree-building methods. Firstly, maximum parsimony, and maximum parsimony with insertions and deletions coded as a 5$^{th}$ character state, analyses were performed in R using the *phangorn* package [9]. Subsequently, maximum likelihood analysis was performed using the software raxmlGUI 2.0, with parameters set to default [10]. Finally, alignment-free k-mer with jack-knifing was performed in R [11]. Code is accessible at GitHub (link will be updated upon acceptance to journal).

Recently described *Cladocopium* species [7] were consistently monophyletic across tree-building methods, with *C. madreporum* and *C. patulum* forming well-supported sister clades (Figure S2). Monophyly of C3f and C3k consistent with ITS2 profile distinction was recovered across analyses, although placement varied (Figure S2), hence C3f and C3k were considered distinct lineages.

**Linear deconvolution**

Linear deconvolution analysis was carried to test for instances where intra-sample co-occurrence of C3f and C3k may have been masked in SymPortal ITS2 profile output. Reference C3f and C3k profiles were generated from samples where the presence of C3f or C3k, detected by ITS2 sequencing, was validated by psbA$^{ncr}$ sequence analysis (n = 8 each) (see above). The frequency of occurrence for each sequence across samples was determined, with sequences appearing in at least 7 samples retained. For each of C3f and C3k, the mean relative abundance and associated standard deviation of each sequence across samples was then generated to form reference profiles (Tables S1 and S2). Only samples where greater than 95% of ITS2 sequences were attributable to *Cladocopium* and total read depth exceeded 5000 were used for reference profile generation. Subsequently, all samples where ITS2 type profiles indicated the presence of *Cladocopium* (C3k or C3f) (n = 195) were deconvoluted against C3f and C3k reference profiles using the *deconvolute* function in the deconvR package [12], following a non-negative least squares model.

Five samples were identified with intermediate contributions from C3f and C3k reference profiles (0.4-0.6) (Table S3, Figure S1). ITS2 type profile out identified these samples as either C3/C3f-C3b-C3.10-C3fl-C3gj-C3ae-C21 (2/5 samples) or C3/C3k-C50a-C3.10-C3gj-C3b (3/5 samples) (Figure S1). The full ITS2 sequence composition across samples is provided as Table S4.

**Table S1.** C3f reference profile derived from ITS2 sequences identified in psbA$^{ncr}$-validated C3f samples

| Name | Sequence | Mean proportion | SD | Number of samples where present |
|---|---|---|---|---|
| C3 | AATGGCCTCCTGAACGTGCGTTGCACTCTTGGGATTTCCTGAGAGTATGTCTGCTTCAGTGCTTAACTTGCCCCAACTTTGCAAGCAGGATGTGTTTCTGCCTTGCGTTCTTATGAGCTATTGCCCTCTGAGCCAATGGCTTGTTAATTGCTTGGTTCTTGCAAAATGCTTTGCGCGCTGTTATTCAAGTTTCTACCTTCGTGGTTTTACTTGAGTGACGCTGCTCATGCTTGCAACCGCTGGGATGCAGGTGCATGCCTCTA | 0.41050945 | 0.021385 | 8 |
| C3f | AGTGGCCTCCTGAATGTGCGTTGCACTCTTGGGATTTCCTGAGAGTATGTCTGCTTCAGTGCTTAACTTGCCCCAACTTTGCAAGCAGGATGTGTTTCTGCCTTGCGTTCGTATGAGCTGTTGCCCTCTGAGCCAATGGCTTGTTAATTGCTTGGTTCTTGCAAAATGCTTTGCGCGCTGTTATTCAAGTTTCTGCCTTCGCGGTTTTACTTGAGTGATGCTGCTCATGCTTGCAACCGCTGGGATGCAGGTGCATGCCTCTA | 0.11069602 | 0.045686 | 8 |
| C3b | AATGGCCTCCTGAACGTGCGTTGCACTCTTGGGATTTCCTGAGAGTATGTCTGCTTCAGTGCTTAACTTGCCCAACTTTGCAAGCAGGATGTGTTTCTGCCTTGCGTTCTTATGAGCTATTGCCCTCTGAGCCAATGGCTTGTTAATTGCTTGGTTCTTGCAAAATGCTTTGCGCGCTGTTATTCAAGTTTCTACCTTCGTGGTTTTACTTGAGTGACGCTGCTCATGCTTGCAACCGCTGGGATGCAGGTGCATGCCTCTA | 0.057307026 | 0.01062 | 8 |
| C3.10 | AATGGCCTCCTGAACGTGCGTTGCACTCTTGGGATTTCCTGAGAGTATGTCTGCTTCAGTGCTTAACTTGCCCCAACTTTGCAAGCAGGATGTGTTTCTGCCTTGCGTTCTTATGAGCTATTGCCCTCTGAGCCAATGGCTTGTTAATTGCTTGGTTCTTGCAAAATGCTTTGCGCGCTGTTATTCAAGTTTCTACCTTCGTGGTTTTACTTGAGTGACGCTGCTCATGCTTGCAACTGCTGGGATGCAGGTGCATGCCTCTA | 0.052945127 | 0.010497 | 8 |
| C3gj | AATGGCCTCCTGAACGTGCGTTGCACTCTTGGGATTTCCTGAGAGTATGTCTATTTCAGTGCTTAACTTGCCCCAACTTTGCAAGCAGGATGTGTTTCTGCCTTGCGTTCTTATGAGCTATTGCCCTCTGAGCCAATGGCTTGTTAATTGCTTGGTTCTTGCAAAATGCTTTGCGCGCTGTTATTCAAGTTTCTACCTTCGTGGTTTTACTTGAGTGACGCTGCTCATGCTTGCAACCGCTGGGATGCAGGTGTATGCCTCTA | 0.048015562 | 0.009046 | 8 |
| C3fl | AATGGCCTCCTGAATGTGCGTTGCACTCTTGGGATTTCCTGAGAGTATGTCTGCTTCAGTGCTTAACTTGACCCAACTTTGCAAGCAGGATGTGTTTCTGCCTTGCGTTCGTATGAGCTGTTGCCCTCTGAGCCAATGGCTTGTTAATTGCTTGGTTCTTGCAAAATGCTTTGCGTGCTGTTATTCAAGTTTCTACCTTCGCGGTTTTACTTGAGTGATGCTGCTCATGCTTGCAACCGCTGGGATGCAGGTGCATGCCTCTA | 0.038290275 | 0.02614 | 8 |
| C21 | AATGGCCTCCTGAACGTGCGTTGCACTCTTGGGATTTCCTGAGAGTATGTCTGCTTCAGTGCTTAACTTGCCCCAACTTTGCAAGCATTTCTGCCTTGCGTTCTTATGAGCTATTGCCCTCTGAGCCAATGGCTTGTTAATTGCTTGGTTCTTGCAAAATGCTTTGCGCGCTGTTATTCAAGTTTCTACCTTCGTGGTTTTACTTGAGTGACGCTGCTCATGCTTGCAACCGCTGGGATGCAGGTGCATGCCTCTA | 0.036363727 | 0.015018 | 8 |
| C3up | AATGGCCTCCTGAACGTGCGTTGCACTTTTGGGATTTCCTGAGAGTATGTCTGCTTCAGTGCTTAACTTGCCCCAACTTTGCAAGCAGGATGTGTTTCTGCCTTGCGTTCTTATGAGCTATTGCCCTCTGAGCCAATGGCTTGTTAATTGCTTGGTTCTTGCAAAATGCTTTGCGCGCTGTTATTCAAGTTTCTACCTTCGTGGTTTTACTTGAGTGACGCTGCTCATGCTTGCAACCGCTGGGATGCAGGTGCATGCCTCTA | 0.034163872 | 0.009569 | 8 |
| C3ae | AATGGCCTCCTGAATGTGCGTTGCACTCTTGGGATTTCCTGAGAGTATGTCTGCTTCAGTGCTTAACTTGCCCCAACTTTGCAAGCAGGATGTGTTTCTGCCTTGCGTTCGTATGAGCTGTTGCCCTCTGAGCCAATGGCTTGTTAATTGCTTGGTTCTTGCAAAATGCTTTGCGTGCTGTTATTCAAGTTTCTACCTTCGCGGTTTTACTTGAGTGATGCTGCTCATGCTTGCAACCGCTGGGATGCAGGTGCATGCCTCTA | 0.026686863 | 0.014529 | 8 |
| 37348_C | AATGGCCTCCTGAACGTGCGTTGCACTCTTGGGATTTCCTGAGAGTATGTCTATTTCAGTGCTTAACTTGCCCCAACTTTGCAAGCAGGATGTGTTTCTGCCTTGCGTTCTTATGAGCTATTGCCCTCTGAGCCAATGGCTTGTTAATTGCTTGGTTCTTGCAAAATGCTTTGCGCGCTGTTATTCAAGGTTCTACCTTCGTGGTTTTACTTGAGTGACGCTGCTCATGCTTGCAACCGCTGGGATGCAGGTGTATGCCTCTA | 0.025907548 | 0.00669 | 8 |

| | | | | |
|---|---|---|---|---|
| C50a | AATGGCCTCCTGAATGTGCGTTGCACTCTTGGGATTTCCTGAGAGTATGTCTGCTTCAGTGCTTAACTTGCCCCAACTTTGCAAGCAGGATGTGTTTCTGCCTTGCGTTCGTATGAGCTATTGCCCTCTGAGCCAATGGCTTGTTAATTGCTTGGTTCTTGCAAAATGCTTTGCGCGCTGTTATTCAAGTTTCTACCTTCGTGGTTTTACTTGAGTGACGCTGCTCATGCTTGCAACCGCTGGGATGCAGGTGCATGCCTCTA | 0.025676175 | 0.007604 | 8 |
| 3638_C | AATGGCCTCCTGAACGTGCGTTGCACTCTTGGGATTTCCTGAGAGTATGTCTGCTTCAGTGCTTAACTTGCCCCAACTTTGCAAGCAGGATGTGTTTCTGCCTTGCGTTCTTATGAGCTATTGCCCTCTGAGCCAATGGCTTGTTAATTGCTTGGTTCTTGCAAAATGCTTTGCGCGCTGTTATTCAAGTTTCTACCTTCGTGGTTTTACTTGAGTGACGCTGCTCATGCTTGCAACCGCTGGGATGCAGGTGCGTGCCTCTA | 0.02497325 | 0.007708 | 8 |
| C3.8 | AATGGCCTCCTGAACGTGCGTTGCACTCTTGGGATTTCCTGAGAGTATGTCTGCTTCAGTGCTTAACTTGCCCCAACTTTGCAAGCAGGATGTGTTTCTGCCTTGCGTTCTTATGAGCTATTGCCCTCTGAGCCAATGGCTTGTTAATTGCTTGGTTCTTGCAAAATGCTTTGCGCGCTGTTATTCAAGTTTCTATCTTCGTGGTTTTACTTGAGTGACGCTGCTCATGCTTGCAACCGCTGGGATGCAGGTGCATGCCTCTA | 0.024803887 | 0.007029 | 8 |
| C3zl | AATGGCCTCCTGAACGTGCGTTGCACTCTTGGGATTTCCTGAGAGTATGTCTACTTCAGTGCTTAACTTGCCCCAACTTTGCAAGCAGGATGTGCTTCTGCCTTGCGTTCTTATGAGCTATTGCCCTCTGAGCCAATGGCTTGTTAATGGCTTGGTTATTGCAAAATGCTTTGCGCGCTGTTATTCAAGTTTCTACCTTCGTGGTTTTACTTGAGTGACGCTGCTCATGCTTGCAACCGCTGGGATGCAGGTGCATGCCTCTA | 0.024323261 | 0.008074 | 8 |
| 21810_C | AATGGCCTCCTGAACGTGCGTTGCACTCTTGGGATTTCCTGAGAGTATGTCTGCTTCAGTGCTTAACTTGCCCCAACTTTGCAAGCAGGATGTGTTTCTGCCTTGCGTTCTTATGAGCTATTGCCCTCTGAGCCAATGGCTTGTTAATTGCTTGGTTCTTGCAAAATGCTTTGCGCGCTGTTATTCAAGTTTCTACCTTCGTGGTTTTACTTGAGTGACGCTGCTCATGCTTGCAACCTCTGGGATGCAGGTGCATGCCTCTA | 0.019964138 | 0.005967 | 8 |
| C21cf | AATGGCTTCGTGAACGTGCGTTGCACTCTTGGGATTTCCTGAGAGTATGTCTGCTTCAGTGCTTAACTTGCCCCAACTTTGCAAGCATTTCTGCCTTGCGTTCTTATGAGCTATTGCCCTCTGAGCCAATGGCTTGTTAATTGCTTGGTTCTTGCAAAATGCTTTGCGCGCTGTTATTCAAGTTTCTACCTTCGTGGTTTTACTTGAGTGACGCTGCTCATGCTTGCAACCGCTGGGATGCAGGTGCATGCCTCTA | 0.018080763 | 0.007695 | 8 |
| C3nf | AATGGCCTCCTGAACGTGCGTTGCACTCTTGGGATTTCCTGAGAGTATGTCTGCTTCAGTGCTTAACTTGCCCCAACTTTGCAAGCAGGATGTGCTTCTGCCTTGCGTTCTTATGAGCTATTGCCCTCTGAGCCAATGGCTTGTTAATGGCTTGGTTCTTGCAAAATGCTTTGCGCGCTGTTATTCAAGTTTCTACCTTCGTGGTTTTACTTGAGTGACGCTGCTCATGCTTGCAACCGCTGGGATGCAGGTGCATGCCTCTA | 0.011883363 | 0.004121 | 8 |
| C3gl | AATGGCCTCCTGAACGTGCGTTGCACTCTTGCGATTTCCTGAGAGTATGTCTGCTTCAGTGCTTAACTTGCCCCAACTTTGCAAGCAGGATGTGTTTCTGCCTTGCGTTCTTATGAGCTATTGCCCTCTGAGCCAATGGCTTGTTAATTGCTTGGTTCTTGCAAAATGCTTTGCGCGCTGTTATTCAAGTTTCTACCTTCGTGGTTTTACTTGAGTGACGCTGCTCATGCTTGCAACCGCTGGGATGCAGGTGCATGCCTCTA | 0.009409693 | 0.003224 | 8 |

**Table S2.** C3k reference profile derived from ITS2 sequences identified in psbA[ncr]-validated C3k samples

| **Name** | **Sequence** | **Average proportion** | **SD** | **Number of samples where present** |
|---|---|---|---|---|
| C3 | AATGGCCTCCTGAACGTGCGTTGCACTCTTGGGATTTCCTGAGAGTATGTCTGCTTCAGTGCTTAACTTGCCCCAACTTTGCAAGCAGGATGTGTTTCTGCCTTGCGTTCTTATGAGCTATTGCCCTCTGAGCCAATGGCTTGTTAATTGCTTGGTTCTTGCAAAATGCTTTGCGCGCTGTTATTCAAGTTTCTACCTTCGTGGTTTTACTTGAGTGACGCTGCTCATGCTTGCAACCGCTGGGATGCAGGTGCATGCCTCTA | 0.480787678 | 0.022118 | 8 |
| C3k | AATGGCCTCCTGAATGTGCGTTGCACTCTTGGGATTTCCTGAGAGTATGTCTGCTTCAGTGCTTAACTTGCCCCAACTTTGCAAGCAGGATGTGTTTCTGCCTTGCGTTCGTATGAGCTGTTGCCCTCTGAGCCAATGGCTTGTTAATTGCTTGGTTCTTGCAAAATGCTTTGCGCGCTGTTATTCAAGTTTCTACCTTCGTGGTTTTACTTGAGTGATGCTGCTCATGCTTGCAACCGCTGGGATGCAGGTGCATGCCTCTA | 0.240059495 | 0.029409 | 8 |

| | | | | |
|---|---|---|---|---|
| C50a | AATGGCCTCCTGAATGTGCGTTGCACTCTTGGGATTTCCTGAGAGTATGTCTGCTTCAGTGCTTAACTTGCCCCAACTTTGCAAGCAGGATGTGTTTCTGCCTTGCGTTCGTATGAGCTATTGCCCTCTGAGCCAATGGCTTGTTAAATTGCTTGGTTCTTGCAAAATGCTTTGCGCGCTGTTATTCAAGTTTCTACCTTCGTGGTTTTACTTGAGTGACGCTGCTCATGCTTGCAACCGCTGGGATGCAGGTGCATGCCTCTA | 0.065919916 | 0.009372 | 8 |
| C3.10 | AATGGCCTCCTGAACGTGCGTTGCACTCTTGGGATTTCCTGAGAGTATGTCTGCTTCAGTGCTTAACTTGCCCCAACTTTGCAAGCAGGATGTGTTTCTGCCTTGCGTTCTTATGAGCTATTGCCCTCTGAGCCAATGGCTTGTTAATTGCTTGGTTCTTGCAAAATGCTTTGCGCGCTGTTATTCAAGTTTCTACCTTCGTGGTTTTACTTGAGTGACGCTGCTCATGCTTGCAACTGCTGGGATGCAGGTGCATGCCTCTA | 0.055948346 | 0.015347 | 8 |
| C3gj | AATGGCCTCCTGAACGTGCGTTGCACTCTTGGGATTTCCTGAGAGTATGTCTATTTCAGTGCTTAACTTGCCCCAACTTTGCAAGCAGGATGTGTTTCTGCCTTGCGTTCTTATGAGCTATTGCCCTCTGAGCCAATGGCTTGTTAATTGCTTGGTTCTTGCAAAATGCTTTGCGCGCTGTTATTCAAGTTTCTACCTTCGTGGTTTTACTTGAGTGACGCTGCTCATGCTTGCAACCGCTGGGATGCAGGTGTATGCCTCTA | 0.037761194 | 0.010973 | 8 |
| C3b | AATGGCCTCCTGAACGTGCGTTGCACTCTTGGGATTTCCTGAGAGTATGTCTGCTTCAGTGCTTAACTTGCCCAACTTTGCAAGCAGGATGTGTTTCTGCCTTGCGTTCTTATGAGCTATTGCCCTCTGAGCCAATGGCTTGTTAATTGCTTGGTTCTTGCAAAATGCTTTGCGCGCTGTTATTCAAGTTTCTACCTTCGTGGTTTTACTTGAGTGACGCTGCTCATGCTTGCAACCGCTGGGATGCAGGTGCATGCCTCTA | 0.0304703 | 0.007412 | 8 |
| C21 | AATGGCCTCCTGAACGTGCGTTGCACTCTTGGGATTTCCTGAGAGTATGTCTGCTTCAGTGCTTAACTTGCCCCAACTTTGCAAGCATTTCTGCCTTGCGTTCTTATGAGCTATTGCCCTCTGAGCCAATGGCTTGTTAATTGCTTGGTTCTTGCAAAATGCTTTGCGCGCTGTTATTCAAGTTTCTACCTTCGTGGTTTTACTTGAGTGACGCTGCTCATGCTTGCAACCGCTGGGATGCAGGTGCATGCCTCTA | 0.02392117 | 0.00411 | 8 |
| 21204_C | AATGGCCTCCTGAACGTGCGTTGCACTCTTGGGATTTCCTGAGAGTATGTCTGCTTCAGTGCTTAACTTGCCCCAACTTTGCAAGCAGGATGTGTTTCTCCCTTGCGTTCTTATGAGCTATTGCCCTCTGAGCCAATGGCTTGTTAATTGCTTGGTTCTTGCAAAATGCTTTGCGCGCTGTTATTCAAGTTTCTACCTTCGTGGTTTTACTTGAGTGACGCTGCTCATGCTTGCAACCGCTGGGATGCAGGTGCATGCCTCTA | 0.023370744 | 0.008668 | 8 |
| C21cf | AATGGCTTCGTGAACGTGCGTTGCACTCTTGGGATTTCCTGAGAGTATGTCTGCTTCAGTGCTTAACTTGCCCCAACTTTGCAAGCATTTCTGCCTTGCGTTCTTATGAGCTATTGCCCTCTGAGCCAATGGCTTGTTAATTGCTTGGTTCTTGCAAAATGCTTTGCGCGCTGTTATTCAAGTTTCTACCTTCGTGGTTTTACTTGAGTGACGCTGCTCATGCTTGCAACCGCTGGGATGCAGGTGCATGCCTCTA | 0.021986548 | 0.007529 | 8 |
| C3cz | AATGGCCTCCTGAATGTGCGTTGCACTCTTGGGATTTCCTGAGAGTATGTCTGCTTCAGTGCTTAACTTGCCCCAACTTTGCAAGCAGGATGTGTTTCTGCCTTGCGTTCGTATGAGCTGTTGCCCTCTGAGCCAATGGCTTGTTAATTGCTTGGTTCTTGCAAAATGCTTTGTGCGCTGTTATTCAAGTTTCTACCTTCGTGGTTTTACTTGAGTGATGCTGCTCATGCTTGCAACCGCTGGGATGCAGGTGCATGCCTCTA | 0.009984845 | 0.001712 | 8 |
| 49135_C | AATGGCCTCCTGAACGTGCGTTGCACTCTTGGGATTTCCTGAGAGTATGTCTGCTTCAGTGCTTAACTTGCCCCAACTTTGCAAGCAGGATGTGTTTCTGCCTTGCGTTCTATGAGCTATTGCCCTCTGAGCCAATGGCTTGTTAATTGCTTGGTTCTTGCAAAATGCTTTGCGCGCTGTTATTCAAGTTTCTACCTTCGTGGTTTTACTTGAGTGACACTGCTCATGCTTGCAACCGCTGGGATGCAGGTGCATGCCTCTA | 0.009789763 | 0.002857 | 8 |

**Table S3.** Non-negative least squares (nnls) deconvolution results for samples where *Cladocopium-Cladocopium* co-occurrence was likely masked, showing estimated proportions attributable to reference C3f and C3k profiles and associated R-squared (rsq) values.

| Sample ID | Proportion C3f | Proportion C3k | rsq | Identified ITS2 type *Cladocopium* profile |
|---|---|---|---|---|
| CC5 | 0.678498959 | 0.321501041 | 0.24067752 | C3/C3f-C3b-C3.10-C3fl-C3gj-C3ae-C21 |
| L4 | 0.623183300 | 0.376816700 | 0.45794942 | C3/C3f-C3b-C3.10-C3fl-C3gj-C3ae-C21 |
| Y1 | 0.432453490 | 0.567546510 | 0.06426394 | C3/C3k-C50a-C3.10-C3gj-C3b |
| J2 | 0.419816623 | 0.580183377 | 0.33031265 | C3/C3k-C50a-C3.10-C3gj-C3b |
| EE2 | 0.414827142 | 0.585172858 | 0.14330646 | C3/C3k-C50a-C3.10-C3gj-C3b |

**Table S4.** Relative abundance of each ITS2 sequence across samples where *Cladocopium-Cladocopium* co-occurrence was likely masked

| name | sequence | CC5 | L4 | J2 | EE2 | Y1 |
|---|---|---|---|---|---|---|
| C3 | AATGGCCTCCTGAACGTGCGTTGCACTCTTGGGATTTCCTGAGAGTATGTCTGCTTC<br>AGTGCTTAACTTGCCCCAACTTTGCAAGCAGGATGTGTTTCTGCCTTGCGTTCTTAT<br>GAGCTATTGCCCTCTGAGCCAATGGCTTGTTAATTGCTTGGTTCTTGCAAAATGCTTT<br>GCGCGCTGTTATTCAAGTTTCTACCTTCGTGGTTTTACTTGAGTGACGCTGCTCATGC<br>TTGCAACCGCTGGGATGCAGGTGCATGCCTCTA | 0.379787 | 0.324565 | 0.42542 | 0.383325 | 0.379168 |
| C3k | AATGGCCTCCTGAATGTGCGTTGCACTCTTGGGATTTCCTGAGAGTATGTCTGCTTC<br>AGTGCTTAACTTGCCCCAACTTTGCAAGCAGGATGTGTTTCTGCCTTGCGTTCGTAT<br>GAGCTGTTGCCCTCTGAGCCAATGGCTTGTTAATTGCTTGGTTCTTGCAAAATGCTTT<br>GCGCGCTGTTATTCAAGTTTCTACCTTCGTGGTTTTACTTGAGTGATGCTGCTCATGC<br>TTGCAACCGCTGGGATGCAGGTGCATGCCTCTA | 0.084636 | 0.075819 | 0.124505 | 0.13364 | 0.135886 |
| C3f | AGTGGCCTCCTGAATGTGCGTTGCACTCTTGGGATTTCCTGAGAGTATGTCTGCTTC<br>AGTGCTTAACTTGCCCCAACTTTGCAAGCAGGATGTGTTTCTGCCTTGCGTTCGTAT<br>GAGCTGTTGCCCTCTGAGCCAATGGCTTGTTAATTGCTTGGTTCTTGCAAAATGCTTT<br>GCGCGCTGTTATTCAAGTTTCTGCCTTCGCGGTTTTACTTGAGTGATGCTGCTCATGC<br>TTGCAACCGCTGGGATGCAGGTGCATGCCTCTA | 0.114775 | 0.071197 | 0.028704 | 0.07032 | 0.084057 |
| C3.10 | AATGGCCTCCTGAACGTGCGTTGCACTCTTGGGATTTCCTGAGAGTATGTCTGCTTC<br>AGTGCTTAACTTGCCCCAACTTTGCAAGCAGGATGTGTTTCTGCCTTGCGTTCTTAT<br>GAGCTATTGCCCTCTGAGCCAATGGCTTGTTAATTGCTTGGTTCTTGCAAAATGCTTT<br>GCGCGCTGTTATTCAAGTTTCTACCTTCGTGGTTTTACTTGAGTGACGCTGCTCATGC<br>TTGCAACTGCTGGGATGCAGGTGCATGCCTCTA | 0.034897 | 0.038696 | 0.041669 | 0.057518 | 0.047181 |
| C3b | AATGGCCTCCTGAACGTGCGTTGCACTCTTGGGATTTCCTGAGAGTATGTCTGCTTC<br>AGTGCTTAACTTGCCCAACTTTGCAAGCAGGATGTGTTTCTGCCTTGCGTTCTTATG<br>AGCTATTGCCCTCTGAGCCAATGGCTTGTTAATTGCTTGGTTCTTGCAAAATGCTTTG<br>CGCGCTGTTATTCAAGTTTCTACCTTCGTGGTTTTACTTGAGTGACGCTGCTCATGCT<br>TGCAACCGCTGGGATGCAGGTGCATGCCTCTA | 0.019601 | 0.031222 | 0.047764 | 0.027813 | 0.018388 |
| C50a | AATGGCCTCCTGAATGTGCGTTGCACTCTTGGGATTTCCTGAGAGTATGTCTGCTTC<br>AGTGCTTAACTTGCCCCAACTTTGCAAGCAGGATGTGTTTCTGCCTTGCGTTCGTAT<br>GAGCTATTGCCCTCTGAGCCAATGGCTTGTTAATTGCTTGGTTCTTGCAAAATGCTTT<br>GCGCGCTGTTATTCAAGTTTCTACCTTCGTGGTTTTACTTGAGTGACGCTGCTCATGC<br>TTGCAACCGCTGGGATGCAGGTGCATGCCTCTA | 0.03399 | 0.03255 | 0.046172 | 0.038156 | 0.041625 |
| C3gj | AATGGCCTCCTGAACGTGCGTTGCACTCTTGGGATTTCCTGAGAGTATGTCTATTTCA<br>GTGCTTAACTTGCCCCAACTTTGCAAGCAGGATGTGTTTCTGCCTTGCGTTCTTATG<br>AGCTATTGCCCTCTGAGCCAATGGCTTGTTAATTGCTTGGTTCTTGCAAAATGCTTTG<br>CGCGCTGTTATTCAAGTTTCTACCTTCGTGGTTTTACTTGAGTGACGCTGCTCATGCT<br>TGCAACCGCTGGGATGCAGGTGTATGCCTCTA | 0.021754 | 0.024929 | 0.032025 | 0.037462 | 0.017781 |
| C21 | AATGGCCTCCTGAACGTGCGTTGCACTCTTGGGATTTCCTGAGAGTATGTCTGCTTC<br>AGTGCTTAACTTGCCCCAACTTTGCAAGCATTTCTGCCTTGCGTTCTTATGAGCTATT<br>GCCCTCTGAGCCAATGGCTTGTTAATTGCTTGGTTCTTGCAAAATGCTTTGCGCGCT<br>GTTATTCAAGTTTCTACCTTCGTGGTTTTACTTGAGTGACGCTGCTCATGCTTGCAAC<br>CGCTGGGATGCAGGTGCATGCCTCTA | 0.037616 | 0.025617 | 0.022608 | 0.029831 | 0.017175 |
| C3fl | AATGGCCTCCTGAATGTGCGTTGCACTCTTGGGATTTCCTGAGAGTATGTCTGCTTC<br>AGTGCTTAACTTGACCCAACTTTGCAAGCAGGATGTGTTTCTGCCTTGCGTTCGTAT<br>GAGCTGTTGCCCTCTGAGCCAATGGCTTGTTAATTGCTTGGTTCTTGCAAAATGCTTT<br>GCGTGCTGTTATTCAAGTTTCTACCTTCGCGGTTTTACTTGAGTGATGCTGCTCATGC<br>TTGCAACCGCTGGGATGCAGGTGCATGCCTCTA | 0.008384 | 0.015046 | 0.011827 | NA | 0.008386 |
| C21cf | AATGGCTTCGTGAACGTGCGTTGCACTCTTGGGATTTCCTGAGAGTATGTCTGCTTC<br>AGTGCTTAACTTGCCCCAACTTTGCAAGCATTTCTGCCTTGCGTTCTTATGAGCTATT<br>GCCCTCTGAGCCAATGGCTTGTTAATTGCTTGGTTCTTGCAAAATGCTTTGCGCGCT<br>GTTATTCAAGTTTCTACCTTCGTGGTTTTACTTGAGTGACGCTGCTCATGCTTGCAAC<br>CGCTGGGATGCAGGTGCATGCCTCTA | 0.014276 | 0.019028 | 0.022335 | 0.016524 | 0.013033 |
| C3ae | AATGGCCTCCTGAATGTGCGTTGCACTCTTGGGATTTCCTGAGAGTATGTCTGCTTC<br>AGTGCTTAACTTGCCCCAACTTTGCAAGCAGGATGTGTTTCTGCCTTGCGTTCGTAT<br>GAGCTGTTGCCCTCTGAGCCAATGGCTTGTTAATTGCTTGGTTCTTGCAAAATGCTTT<br>GCGTGCTGTTATTCAAGTTTCTACCTTCGCGGTTTTACTTGAGTGATGCTGCTCATGC<br>TTGCAACCGCTGGGATGCAGGTGCATGCCTCTA | 0.044414 | 0.014554 | 0.008598 | 0.027434 | 0.032835 |
| C3up | AATGGCCTCCTGAACGTGCGTTGCACTTTTGGGATTTCCTGAGAGTATGTCTGCTTC<br>AGTGCTTAACTTGCCCCAACTTTGCAAGCAGGATGTGTTTCTGCCTTGCGTTCTTAT<br>GAGCTATTGCCCTCTGAGCCAATGGCTTGTTAATTGCTTGGTTCTTGCAAAATGCTTT<br>GCGCGCTGTTATTCAAGTTTCTACCTTCGTGGTTTTACTTGAGTGACGCTGCTCATGC<br>TTGCAACCGCTGGGATGCAGGTGCATGCCTCTA | 0.01235 | 0.012636 | 0.015921 | 0.006622 | 0.011922 |

| | | | | | | |
|---|---|---|---|---|---|---|
| 21810_C | AATGGCCTCCTGAACGTGCGTTGCACTCTTGGGATTTCCTGAGAGTATGTCTGCTTCAGTGCTTAACTTGCCCCAACTTTGCAAGCAGGATGTGTTTCTGCCTTGCGTTCTTATGAGCTATTGCCCTCTGAGCCAATGGCTTGTTAATTGCTTGGTTCTTGCAAAATGCTTTGCGCGCTGTTATTCAAGTTTCTACCTTCGTGGTTTTACTTGAGTGACGCTGCTCATGCTTGCAACCTCTGGGATGCAGGTGCATGCCTCTA | 0.013596 | 0.013915 | 0.015694 | 0.009523 | 0.01475 |
| 3638_C | AATGGCCTCCTGAACGTGCGTTGCACTCTTGGGATTTCCTGAGAGTATGTCTGCTTCAGTGCTTAACTTGCCCCAACTTTGCAAGCAGGATGTGTTTCTGCCTTGCGTTCTTATGAGCTATTGCCCTCTGAGCCAATGGCTTGTTAATTGCTTGGTTCTTGCAAAATGCTTTGCGCGCTGTTATTCAAGTTTCTACCTTCGTGGTTTTACTTGAGTGACGCTGCTCATGCTTGCAACCGCTGGGATGCAGGTGCGTGCCTCTA | 0.010197 | 0.010621 | 0.012146 | NA | 0.009901 |
| 21204_C | AATGGCCTCCTGAACGTGCGTTGCACTCTTGGGATTTCCTGAGAGTATGTCTGCTTCAGTGCTTAACTTGCCCCAACTTTGCAAGCAGGATGTGTTTCTCCCTTGCGTTCTTATGAGCTATTGCCCTCTGAGCCAATGGCTTGTTAATTGCTTGGTTCTTGCAAAATGCTTTGCGCGCTGTTATTCAAGTTTCTACCTTCGTGGTTTTACTTGAGTGACGCTGCTCATGCTTGCAACCGCTGGGATGCAGGTGCATGCCTCTA | 0.007365 | 0.008555 | 0.006869 | 0.012614 | 0.012023 |
| C3nf | AATGGCCTCCTGAACGTGCGTTGCACTCTTGGGATTTCCTGAGAGTATGTCTGCTTCAGTGCTTAACTTGCCCCAACTTTGCAAGCAGGATGTGCTTCTGCCTTGCGTTCTTATGAGCTATTGCCCTCTGAGCCAATGGCTTGTTAATGGCTTGGTTCTTGCAAAATGCTTTGCGCGCTGTTATTCAAGTTTCTACCTTCGTGGTTTTACTTGAGTGACGCTGCTCATGCTTGCAACCGCTGGGATGCAGGTGCATGCCTCTA | 0.01201 | 0.010719 | 0.010417 | 0.008262 | 0.007779 |
| C3gl | AATGGCCTCCTGAACGTGCGTTGCACTCTTGCGATTTCCTGAGAGTATGTCTGCTTCAGTGCTTAACTTGCCCCAACTTTGCAAGCAGGATGTGTTTCTGCCTTGCGTTCTTATGAGCTATTGCCCTCTGAGCCAATGGCTTGTTAATTGCTTGGTTCTTGCAAAATGCTTTGCGCGCTGTTATTCAAGTTTCTACCTTCGTGGTTTTACTTGAGTGACGCTGCTCATGCTTGCAACCGCTGGGATGCAGGTGCATGCCTCTA | 0.020054 | 0.005999 | 0.006323 | 0.013055 | 0.008183 |
| 183_C | AATGGCCTCCTGAACGTGCGTTGCACTCTTGGGATTTCCTGAGAGTATGTCTGCTTCAGTGCTTAACGTGCCCCAACTTTGCAAGCAGGATGTGTTTCTGCCTTGCGTTCTTATGAGCTATTGCCCTCTGAGCCAATGGCTTGTTAATTGCTTGGTTCTTGCAAAATGCTTTGCGCGCTGTTATTCAAGTTTCTACCTTCGTGGTTTTACTTGAGTGACGCTGCTCATGCTTGCAACCGCTGGGATGCAGGTGCATGCCTCTA | 0.009631 | 0.006539 | 0.00646 | 0.01192 | 0.004748 |
| C3cp | AATGGCCTCCTGAACGTGCGTTGCACTCTTGGGTTTTCCTGAGAGTATGTCTGCTTCAGTGCTTAACTTGCCCCAACTTTGCAAGCAGGATGTGTTTCTGCCTTGCGTTCTTATGAGCTATTGCCCTCTGAGCCAATGGCTTGTTAATTGCTTGGTTCTTGCAAAATGCTTTGCGCGCTGTTATTCAAGTTTCTACCTTCGTGGTTTTACTTGAGTGACGCTGCTCATGCTTGCAACCGCTGGGATGCAGGTGCATGCCTCTA | 0.015749 | 0.007425 | 0.007415 | 0.009397 | 0.009295 |
| C3uq | AATGGCCTCCTGAACGTGCGTTGCACTCTTGGGATTTCCTGAGAGTATGTCTGCTTCAGTGCTTAACTTGCCCAACTTTGCAAGCAGGATGTGTCTCTGCCTTGCGTTCTTATGAGCTATTGCCCTCTGAGCCAATGGCTTGTTAATTGCTTGGTTCTTGCAAAATGCTTTGCGCGCTGTTATTCAAGTTTCTACCTTCGTGGTTTTACTTGAGTGACGCTGCTCATGCTTGCAACCGCTGGGATGCAGGTGCATGCCTCTA | 0.010877 | 0.005507 | 0.008279 | 0.007379 | 0.006365 |
| C3na | AATGGCCTCCTGAACGTGCGTTGCACTCTTGGGATTTCCTGAGAGTATGTCTATTTCAGTGCTTAACTTGCCCCAACTTTGCAAGCAGGATGTGTTTCTGCCTTGCGTTCTTATGAGCTATTACCCTCTGAGCCAATGGCTTGTTAATTGCTTGGTTCTTGCAAAATGCTTTGCGCGCTGTTATTCAAGTTTCTACCTTCGTGGTTTTACTTGAGTGACGCTGCTCATGCTTGCAACCGCTGGGATGCAGGTGTATGCCTCTA | 0.007251 | 0.006834 | 0.005868 | 0.010721 | 0.007577 |
| C3nc | AATGGCCTCCTGAACGTGCGTTGCACTCTTGGGATTTCCTGAGAGTATGTCTGCTTCAGTGCTTAACTTGCCCAACTTTGCAAGCAGGATGTGTTTCTGCCTTGCGCTCTTATGAGCTATTGCCCTCTGAGCCAATGGCTTGTTAATTGCTTGGTTCTTGCAAAATGCTTTGCGCGCTGTTATTCAAGTTTCTACCTTCGTGGTTTTACTTGAGTGACGCTGCTCATGCTTGCAACCGCTGGGATGCAGGTGCATGCCTCTA | 0.010877 | NA | NA | 0.006811 | 0.008891 |
| C3fk | AATGGCCTCCTGAACGTGCGTTGCACTCTTGGGATTTCCTGAGAGTATGTCTGCTTCAGTGCTTAACTTGCCCCAACTTTGCAAGCAGGATGTGTTTCTGCCTTGCGTTCTTATGAGCTATTGCCCTCTGAGCCAATGGCTTGTTAATTGCTTGGTTCTTGCAAAATGCTTTGCGCGCTGTTATTCAAGTTTCTACCTTCGTGGTTTTACTTGAGTGGCGCTGCTCATGCTTGCAACCGCTGGGATGCAGGTGCATGCCTCTA | 0.007591 | 0.005114 | 0.00887 | NA | 0.006971 |
| C3bj | AATGGCCTCCTGAATGTGCGTTGCACTCTTGGGATTTCCTGAGAGTATGTCTGCTTCAGTGCTTAACTTGCCCCAACTTTGCAAGCAGGATGTGTTTCTGCCTTGCGTTCGTATGAGCTGTTGCCCTCTGAGCCAATGGCTTGTTAATTGCTTGGTTCTTGCAAAATGCTTTGCGCGCTGTTATTCAAGTTTCTACCTTCGCGGTTTTACTTGAGTGATGCTGCTCATGCTTGCAACCGCTGGGATGCAGGTGCATGCCTCTA | 0.011897 | NA | NA | NA | 0.007072 |
| 40851_C | AATGGCCTCCTGAACGTGCGTTGCACTCTTGGGATTTCCTGAGAGTATGTCTGCTTCAGTGCTTAACTTGCCCCAACTTTGCAAGCAGGATGTGTTTCTGCCTTGCGTTCTTATGAGCTATTGCCCTCTGAGCCAATGGCTTGTTAATTGCTTGGTTCTTGCAAAATGCTTTGCGCGCTGTTATTCAAGTTTCTACCTTCGTGGTTTTACTTGAGTGGCGCTGCTCATGCTTGCAACCGCTGGGATGCAGATGCATGCCTCTA | 0.008498 | 0.004524 | NA | 0.01501 | 0.005052 |
| 160_C | AATGGCCTCCTGAACGTGCGTTGCACTCTTGGGATTTCCTGAGAGTATGTCTGCTTCAGTGCTTAACTTGCCCAACTTTGCAAGCAGGATGTGTTTCTGCCTTGCGTTCTTATGAGCTATTGCCCTCTGAGCCAATGGCTTGTTAATTGCTTGGTTCTTGCAAAATGCTTTGCGCGCTGTTATTCAAGTTTCTATCTTCGTGGTTTTACTTGAGTGACGCTGCTCATGCTTGCAACCGCTGGGAGCAGGTGCATGCCTCTA | 0.004759 | 0.004376 | NA | 0.008956 | 0.0098 |
| C3fj | AATGGCGTCCTGAACGTGCGTTGCACTCTTGGGATTTCCTGAGAGTATGTCTGCTTCAGTGCTTAACTTGCCCCAACTTTGCAAGCAGGATGTGTTTCTGCCTTGCGTTCTTATGAGCTATTGCCCTCTGAGCCAATGGCTTGTTAATTGCTTGGTTCTTGCAAAATGCTTTGCGCGCTGTTATTCAAGTTTCTACCTTCGTGGTTTTACTTGAGTGACGCTGCTCATGCTTGCAACCGCTGGGATGCAGGTGCATGCCTCTA | 0.006685 | NA | NA | NA | NA |
| C3ur | AATGGCCTCCTGAACGTGCGTTGCACTCTTGGGATTTCCTGAGAGTATGTCTGCTTCAGTGCTTAACTTGCCCCAACTTTGCAAGCAGGATGTGTTTCTGCCTTGCGTTCTTATGAGCTATTGCCCTCTGAGCCAATGGCTTGTTAATTGCTTGGTTCTTGCAAAATGCTTTGACCGCTGTTATTCAAGTTTCTACCTTCGTGGTTTTACTTGAGTGACGCTGCTCATGCTTGCAACCGCTGGGATGCAGGTGCATGCCTCTA | 0.007025 | NA | 0.005504 | NA | NA |
| C15sy | AATGGCCTCCTGAACGTGCGTTGCACTCTTGGGATTTCCTGAGAGTATGTCTGCTTCAGTGCTTAACTTGCCCCAACTTTGCAAGCAGGATGTGTTTCTGCCTTGCGTTCTTATGAGCTATTGCCTTCTGAGCCAATGGCTTGTTAATTGCTTGGTTCTTGCAAAATGCTTTGCGCGCTGTTATTCAAGTTTCTACCTTCGTGGTTTTACTTGAGTGACGCTGCTCATGCTTGCAACCGCTGGGATGCAGGTGCATGCCTCTA | 0.006911 | 0.004917 | NA | NA | 0.009598 |
| C18a | AATGGCCTCCTGAACGTGCGTTGCACTCTTGGGATTTCCTGAGAGTATGTCTGCTTCAGTGCTTAACTTGCCCCAACTTTGCAAGCAGGATGTGTTTCTGCCTTGCGTTCTTATGAGCTATTGCCCTCTGAGCCAATGGCTTGTTAATTGCTTGGTTCTTGCAAAATGCGTTGCGCGCTGTTATTCAAGTTTCTACCTTCGTGGTTTTACTTGAGTGACGCTGCTCATGCTTGCAACCGCTGGGATGCAGGTGCATGCCTCTA | 0.006458 | 0.007474 | 0.005641 | NA | NA |
| C3fc | AATGGCCTCCTGAACGTGCGTTGCACTCTTGGGATTTCCTGAGAGTATGTCTGCTTCAGTGCTTAACTTGCCCCAACTTTGCAAGCAGGATGTGTTTCTGCCTTGCGTTCTTATGAGCTATTGCCCTCTGAGCCAATGGCTTGTTAATTACTTGGTTCTTGCAAAATGCTTTGCGCGCTGTTATTCAAGTTTCTACCTTCGTGGTTTTACTTGAGTGACGCTGCTCATGCTTGCAACCGCTGGGATGCAGGTGCATGCCTCTA | 0.005892 | NA | NA | NA | NA |
| C3nb | AATGGCCTCCTGAACGTGCGTTGCACTCTTGGGATTTCCTGAGAGTATGTCTGCTTCAGTGCTTAACTTGCCCCAACTTTGCAAGCAGGATGTGTTTCTGCCTTGCGTTCTTATGAGCTATTGCCCTCTGAGCCAATGGCTTGTTAATTGCTTGGTTCTTGCAAAATGCTTT | 0.004872 | NA | NA | 0.006054 | NA |

| | | | | | | |
|---|---|---|---|---|---|---|
| | GCGCGCTGTTATTCAAGTTTCTACCTTCGTGGTTTTTTTACTTGAGTGACGCTGCTC<br>ATGCTTGCAACCGCTGGGATGCAGGTGCATGCCTCTA | | | | | |
| C3lu | AATGGCCTCCTGAACGTGCGTTGCACTCTTGGGATTTCCTGAGAGTATGTCTGCTTC<br>AGTGCTTAACTTGCCCCAACTTTGCAAGCAGGATGTGTTTCTGCCTTGCGTTCTTAT<br>GAGCTATTGCCCTCTGAGCCAATGGCTTGTTAATTGCTTGGTTCTTGCAAAATGCTTT<br>GCGCGCTGTTATTCAAGTTTCTACCTTCGTGGTTTTTTTACTTGAGTGACGCTGCTCA<br>TGCTTGCAACCGCTGGGATGCAGGTGCATGCCTCTA | 0.005325 | NA | NA | NA | NA |
| C3zl | AATGGCCTCCTGAACGTGCGTTGCACTCTTGGGATTTCCTGAGAGTATGTCTACTTC<br>AGTGCTTAACTTGCCCCAACTTTGCAAGCAGGATGTGCTTCTGCCTTGCGTTCTTAT<br>GAGCTATTGCCCTCTGAGCCAATGGCTTGTTAATGGCTTGGTTATTGCAAAATGCTTT<br>GCGCGCTGTTATTCAAGTTTCTACCTTCGTGGTTTTACTTGAGTGACGCTGCTCATGC<br>TTGCAACCGCTGGGATGCAGGTGCATGCCTCTA | NA | 0.009293 | 0.012601 | NA | 0.008386 |
| 37348_<br>C | AATGGCCTCCTGAACGTGCGTTGCACTCTTGGGATTTCCTGAGAGTATGTCTATTTCA<br>GTGCTTAACTTGCCCCAACTTTGCAAGCAGGATGTGTTTCTGCCTTGCGTTCTTATG<br>AGCTATTGCCCTCTGAGCCAATGGCTTGTTAATTGCTTGGTTCTTGCAAAATGCTTTG<br>CGCGCTGTTATTCAAGGTTCTACCTTCGTGGTTTTACTTGAGTGACGCTGCTCATGCT<br>TGCAACCGCTGGGATGCAGGTGTATGCCTCTA | NA | 0.007867 | 0.008961 | NA | NA |
| C3.8 | AATGGCCTCCTGAACGTGCGTTGCACTCTTGGGATTTCCTGAGAGTATGTCTGCTTC<br>AGTGCTTAACTTGCCCCAACTTTGCAAGCAGGATGTGTTTCTGCCTTGCGTTCTTAT<br>GAGCTATTGCCCTCTGAGCCAATGGCTTGTTAATTGCTTGGTTCTTGCAAAATGCTTT<br>GCGCGCTGTTATTCAAGTTTCTATCTTCGTGGTTTTACTTGAGTGACGCTGCTCATGC<br>TTGCAACCGCTGGGATGCAGGTGCATGCCTCTA | NA | 0.009883 | 0.009917 | NA | NA |
| C3fm | AATGGCCTCCTGAATGTGCGTTGCACTCTTGGGATTTCCTGAGAGTATGTCTGCTTC<br>AGTGCTTAACTTGACCCAACTTTGCAAGCAGGATGTGTTTCTGCCTTGCGTTCGTAT<br>GAGCTGTTGCCCTCTGAGCCAATGGCTTGTAAATTGCTTGGTTCTTGCAAAATGCTT<br>TGCGTGCTGTTATTCAAGTTTCTACCTTCGCGGTTTTACTTGAGTGATGCTGCTCATG<br>CTTGCAACCGCTGGGATGCAGGTGCATGCCTCTA | NA | 0.008408 | 0.007051 | NA | NA |
| C3ih | AATGGCCTCCTGAACGTGCGTTGCACTCTTGGGATTTCCTGAGAGTATGTCTGCTTC<br>AGTGCTTAACTTGCCCCAACTTTGCAAGCAGGATGTGTTTCTGCCTTGCGTTCTTAT<br>GAGCTATTGCCCTCTGAGCCAATGGCTTGTTAATTGCTTGGTTCTTGCAAAATGCTTT<br>GCGTGCTGTTATTCAAGTTTCTACCTTCGTGGTTTTACTTGAGTGACGCTGCTCATGC<br>TTGCAACCGCTGGGATGCAGGTGCATGCCTCTA | NA | 0.005507 | NA | 0.00864 | 0.009093 |
| C3oc | AATGGCCTCCTGAACGTGCGTTGCACTCCTGGGATTTCCTGAGAGTATGTCTGCTTC<br>AGTGCTTAACTTGCCCCAACTTTGCAAGCAGGATGTGTTTCTGCCTTGCGTTCTTAT<br>GAGCTATTGCCCTCTGAGCCAATGGCTTGTTAATTGCTTGGTTCTTGCAAAATGCTTT<br>GCGCGCTGTTATTCAAGTTTCTACCTTCGTGGTTTTACTTGAGTGACGCTGCTCATGC<br>TTGCAACCGCTGGGATGCAGGTGCATGCCTCTA | NA | 0.004425 | NA | 0.007442 | 0.008689 |
| D1 | AATGGCCCCCTGAACGCGCATTGCACTCTTGGGACTTCCTGAGAGTATGTTTGCTTC<br>AGTGCTTATTTTACCTCCTTGCAAGGTTCTGTCGCAACCTTGTGCCCTGGCCAGCCA<br>TGGGTTAACTTGCCCATGGCTTGCTGAGTAGTGATCTTTTAGAGCAAGCTCTGGCAC<br>GCTGTTGTTTGAGGCAGCCTATATTGAGGCTATTTCAAATGACGTTGCTACAAGCTTG<br>ATGTGTCCTTCTGCGCCGTTGCGCATCCCATA | NA | 0.076605 | 0.000546 | 0.001261 | NA |
| D4 | AATGGCCCCCTGAACGCGCATTGCACTCTTGGGACTTCCTGAGAGTATGTTTGCTTC<br>AGTGCTTATTTTACCTCCTTGCAAGGTTCTGTCGCAACCTTGTGCCCTGGCCAGCCA<br>CGGGTTAACTTGCCCATGGCTTGCTGAGTAGTGATCTTTTAGAGCAAGCTCTGGCAC<br>GCTGTTGTTTGAGGCAGCCTATATTGAGGCTATTTCAAATGACGTTGCTACAAGCTTG<br>ATGTGTCCTTCTGCGCCGTTGCGCATCCCATA | NA | 0.041056 | 0.000864 | 0.000946 | NA |
| D6 | AATGGCCCCCTGAACGCGCATTGCACTCTTGGGACTTCCTGAGAGTATGTTTGCTTC<br>AGTGCTTATTTTACCTCCTTGCAAGGTTCTGTCGCAACCTTGTGCCCTGGCCAGCCA<br>CGCGTTAACTTGCCCATGGCTTGCTGAGTAGTGATCTTTTAGAGCAAGCTCTGGCAC<br>GCTGTTGTTTGAGGCAGCCTATATTGAGGCTATTTCAAATGACGTTGCTACAAGCTTG<br>ATGTGTCCTTCTGCGCCGTTGCGCATCCCATA | NA | 0.018832 | NA | NA | NA |
| D1b | AATGGCCCCCTGAACGCGCATTGCACTCTTGGGACTTCCTGAGAGTATGTTTGCTTC<br>AGTGCTTATTTTACCTCCTTGCAAGGTTCTGTTGCAACCTTGTGCCCTGGCCAGCCAT<br>GGGTTAACTTGCCCATGGCTTGCTGAGTAGTGATCTTTTAGAGCAAGCTCTGGCACG<br>CTGTTGTTTGAGGCAGCCTATATTGAGGCTATTTCAAATGACGTTGCTACAAGCTTGA<br>TGTGTCCTTCTGCGCCGTTGCGCATCCCATA | NA | 0.005999 | NA | 0.000252 | NA |
| D1ab | AATGGCCCCCTGAACGCGCATTGCACTCTTGGGACTTCCTGAGAGTATGTTTGCTTC<br>AGTGCTTATTTTACCTCCTTGCAAGGTTCTGTCGCAACCTTGTGCCCTGGCCAGCCA<br>TGGGTTAACTTGCCCATGGCTTGCTGAGTAGTGATCTTTTAGAGCAAGCTCTGGCAC<br>GCTGTTGTTTGAGGCAGCCTATATTGAGGCTATTTCAAATGACGTTGCTACAGGCTTG<br>ATGTGTCCTCCTGCGCCGTTGCGCATCCCATA | NA | 0.005851 | NA | 0.000315 | NA |
| D1d | AATGGCCCCCTGAACGCGCATTGCACTCTTGGGACTTCCTGAGAGTATGTTTGCTTC<br>AGTGCTTATTTTACCTCCTTGCAAGGTTCTGTCGCAACCTTGTGCCCTGGCCAGCCA<br>TGGGTTAACTTGCCCATGGCTTGCTGAGTAGTGATCTTTTAGAGCAAGCTCTGGCAC<br>GCTGTTGTTTGAGGCAGCCTATATTGAGGCTATTTCAAATGACGTTGCTACAGGCTTG<br>ATGTGTCCTTCTGCGCCGTTGCGCATCCCATA | NA | 0.002753 | NA | 0.000441 | NA |
| D6m | AATGGCCCCCTGAACGCGCATTGCACTCTTGGGACTTCCTGAGAGTATGTTTGCTTC<br>AGTGCTTATTTTAGCTCCTTGCAAGGTTCTGTCGCAACCTTGTGCCCTGGCCAGCCA<br>CGCGTTAACTTGCCCATGGCTTGCTGAGTAGTGATCTTTTAGAGCAAGCTCTGGCAC<br>GCTGTTGTTTGAGGCAGCCTATATTGAGGCTATTTCAAATGACGTTGCTACAAGCTTG<br>ATGTGTCCTTCTGCGCCGTTGCGCATCCCATA | NA | 0.004278 | NA | NA | NA |
| D6c | AATGGCCCCCTGAACGCGCATTGCACTCTTGGGACTTCCTGAGAGTATGTTTGCTTC<br>AGTGCTTATTTTACCTCCTTGCAAGGTTCTGTCGCAACCTTGTGCCCTGGCCAGCCA<br>CGCGTTAACTTGCCCATGGCTTGCTGAGTAGTGATCTTTTAGAGCCAGCTCTGGCAC<br>GCTGTTGTTTGAGGCAGCCTATATTGAGGCTATTTCAAATGACGTTGCTACAAGCTTG<br>ATGTGTCCTTCTGCGCCGTTGCGCATCCCATA | NA | 0.002262 | 0.000227 | NA | NA |
| D2.2 | AATGGCCCCGTGAACGCGCATTGCACTCTTGGGACTTCCTGAGAGTATGTTTGCTTC<br>AGTGCTTATTTTACCTCCTTGCAAGGTTCTGTCGCAACCTTGTGCCCTGGCCAGCCA<br>CGGGTTAACTTGCCCATGGCTTGCTGAGTAGTGATCTTTTAGAGCAAGCTCTGGCAC<br>GCTGTTGTTTGAGGCAGCCTATATTGAGGCTATTTCAAATGACGTTGCTACAAGCTTG<br>ATGTGTCCTTCTGCGCCGTTGCGCATCCCATA | NA | 0.001475 | NA | NA | NA |
| D10d | AATGGCCCCCTGAATGCGCATTGCACTCTTGGGACTTCCTGAGAGTATGTTTACTTC<br>AGTGCTTATTTTACCTCCTTGCAAGGTTCTGTCGCAACCTTGTGCCCTGGCCAGCCA<br>TGGGTTAACTTGCCCATGGCTTGCTGAGTAGTGATCTTTTAGAGCAAGCTCTGGCAC<br>GCTGTTGTTTGAGGCAGCCTATATTGAGGCTATTTCAAATGACGTTGCTACAAGCTTG<br>ATGTGTCCTTCTGCGCCGTTGCGCATCCCATA | NA | 0.001623 | NA | NA | NA |
| D4al | AATGGCCCCCTGAACGCGCATTGCACTCTTGGGACTTCCTGAGAGTATGTTTGCTTC<br>AGTGCTTATTTTACCTCCTTGCAAGGTTTTGTCGCAACCTTGTGCCCTGGCCAGCCA<br>CGGGTTAACTTGCCCATGGCTTGCTGAGTAGTGATCTTTTAGAGCAAGCTCTGGCAC<br>GCTGCTGTTTGAGGCAGCCTATATTGAGGCTATTTCAAATGACGTTGCTACAAGCTT<br>GATGTGTCCTTCTGCGCCGTTGCGCATCCCATA | NA | 0.000885 | NA | NA | NA |
| D4d | AATGGCCCCCTGAACGCGCATTGCACTCTTGGGACTTCCTGAGAGTATGTTTGCTTC<br>AGTGCTTATTTTACCTCCTTGCAAGGTTCTGTCGCAACCTTGTGCCCTGGCCAGCCA<br>CGGGTTAACTTGCCCATGGCTTGCTGAGTAGTGATCTTTTAGAGCAAGCTCTGGCAC<br>GCTGCTGTTTGAGGCAGCCTATATTGAGGCTATTTCAAATGACGTTGCTACAAGCTT<br>GATGTGTCCTTCTGCGCCGTTGCGCATCCCATA | NA | 0.000885 | NA | NA | NA |
| D4am | AATGGCCCCCTGCACGCGCATTGCACTCTTGGGACTTCCTGAGAGTATGTTTGCTTC<br>AGTGCTTATTTTACCTCCTTGCAAGGTTCTGTCGCAACCTTGTGCCCTGGCCAGCCA | NA | 0.001033 | NA | NA | NA |

| | | | | | | |
|---|---|---|---|---|---|---|
| | CGGGTTAACTTGCCCATGGCTTGCTGAGTAGTGATCTTTTAGAGCAAGCTCTGGCAC<br>GCTGTTGTTTGAGGCAGCCTATATTGAGGCTATTTCAAATGACGTTGCTACAAGCTTG<br>ATGTGTCCTTCTGCGCCGTTGCGCATCCCATA | | | | | |
| 914812_D | AATGGCCCCCTGAACGCGCATTGCACTCTTGGGACTTCCTGAGAGTATGTTTGCTTC<br>AGTGCTTATTTTAGCTCCTTGCAAGGTTCTGTCGCAACCTTGTGCCGTGGCCAGCCA<br>CGCGTTAACTTGCCCATGGCTTGCTGAGTAGTGATCTTTTAGAGCAAGCTCTGGCAC<br>GCTGTTGTTTGAGGCAGCCTATATTGAGGCTATTTCAAATGACGTTGCTACAAGCTTG<br>ATGTGTCCTTCTGCGCCGTTGCGCATCCCATA | NA | 0.001033 | NA | NA | NA |
| D1r | AATGGCCTCCTGAACGCGCATTGCACTCTTGGGACTTCCTGAGAGTATGTTTGCTTC<br>AGTGCTTATTTTACCTCCTTGCAAGGTTCTGTCGCAACCTTGTGCCCTGGCCAGCCA<br>TGGGTTAACTTGCCCATGGCTTGCTGAGTAGTGATCTTTTAGAGCAAGCTCTGGCAC<br>GCTGTTGTTTGAGGCAGCCTATATTGAGGCTATTTCAAATGACGTTGCTACAAGCTTG<br>ATGTGTCCTTCTGCGCCGTTGCGCATCCCATA | NA | 0.000983 | NA | NA | NA |
| 3047_D | AATGGCCCCCTGAACGCGCATTGCACTCTTGGGACTTCCTGAGAGTATGTTTGCTTC<br>AGTGCTTATTTTACCTCCTTGCAAGGTTCTGTCGCAACCTTGTGCCCTGGCCAGCCA<br>CGGGTTAACTTGCCCATGGCTTGCTGAGTAGTGATCTTTTAGAGCAAGCTCTGGCAC<br>GCTGTTGTTTGAGGCAGCCTATATTGAGGCTATTTCAAATGACGTTGCTACAAGCTTG<br>ATGTGTCCCTCTGCGCCGTTGCGCATCCCATA | NA | 0.000688 | NA | NA | NA |
| 20890_C | AATGGCCTCCTGAACGTGCGTTGCACTCTTGGGATTTCCTGAGAGTATGTCTGCTAC<br>AGTGCTTAACTTGCCCCAACTTTGCAAGCAGGATGTGTTTCTGCCTTGCGTTCTTAT<br>GAGCTATTGCCCTCTGAGCCAATGGCTTGTTAATTGCTTGGTTCTTGCAAAATGCTTT<br>GCGCGCTGTTATTCAAGTTTCTACCTTCGTGGTTTTACTTGAGTGACGCTGCTCATGC<br>TTGCAACCGCTGGGATGCAGGTGCATGCCTCTA | NA | NA | 0.005732 | 0.005802 | NA |
| C3uo | AATGGCCTCCTGAACGTGCGTTGCACTCTTGGGATTTCCTGAGAGTATGTCTGCTTC<br>AGTGCTTAACTTGCCACAACTTTGCAAGCAGGATGTGTTTCTGCCTTGCGTTCTTAT<br>GAGCTATTGCCCTCTGAGCCAATGGCTTGTTAATTGCTTGGTTCTTGCAAAATGCTTT<br>GCGCGCTGTTATTCAAGTTTCTACCTTCGTGGTTTTACTTGAGTGACGCTGCTCATGC<br>TTGCAACCGCTGGGATGCAGGTGCATGCCTCTA | NA | NA | 0.005095 | NA | NA |
| C3hv | AATGGCCTCCTGAACGTGCGTTGCACTCTTGGGATTTCCTGAGAGTATGTCTGCTTC<br>AGTGCTTAACTTGCCCCAACTTTGCAAGCAGGATGTGTTTCTGCCTTGCGTTCTATG<br>AGCTATTGCCCTCTGAGCCAATGGCTTGTTAATTGCTTGGTTCTTGCAAAATGCTTTG<br>CGCGCTGTTATTCAAGTTTCTACCTTCGTGGTTTTACTTGAGTGACGCTGCTCATGCT<br>TGCAACCGCTGGGATGCAGGTGCATGCCTCTA | NA | NA | 0.006232 | NA | NA |
| C50q | AATGGCCTCCTGAATGTGCGTTGCACTCTTGGGATTTCCTGAGAGTATGTCTGCTTC<br>AGTGCTTAACTTGCCCCAACTTTGCAAGCAGGATGTGTTTCTGCCTTGCGTTCGTAT<br>GAGCTATTGCCCTCTGAGCCAATGGCTTGTTAATTGCTTGGTTCTTGCAAAATGCTTT<br>GCGCGCTGTTATTCAAGTTTCTACCTTCGTGGTTTTACTTGAGTGATGCTGCTCATGC<br>TTGCAACCGCTGGGATGCAGGTGCATGCCTCTA | NA | NA | 0.007551 | NA | 0.007577 |
| C50f | AATGGCCTCCTGAATGTGCGTTGCACTCTTGGGATTTCCTGAGAGTATGTCTGCTTC<br>AGTGCTTAACTTGCCCCAACTTTGCAAGCAGGATGTGTTTCTGCCTTGCGTTCTTAT<br>GAGCTATTGCCCTCTGAGCCAATGGCTTGTTAATTGCTTGGTTCTTGCAAAATGCTTT<br>GCGCGCTGTTATTCAAGTTTCTACCTTCGTGGTTTTACTTGAGTGACGCTGCTCATGC<br>TTGCAACCGCTGGGATGCAGGTGCATGCCTCTA | NA | NA | 0.007733 | 0.00555 | NA |
| C3ng | AATGGCCTCCTGAACGTGCGTTGCACTCTTGGGATTTCCTGAGAGTATGTCTGCTTC<br>AGTGCTTAACTTGCCCCAACTTTGCAAGCAGGATGTGCTTCTGCCTTGCGTTCTTAT<br>GAGCTATTGCCCTCTGAGCCAATGGCTTGTTAATGGCTTGGTTATTGCAAAATGCTTT<br>GCGCGCTGTTATTCAAGTTTCTACCTTCGTGGTTTTACTTGAGTGACGCTGCTCATGC<br>TTGCAACCGCTGGGATGCAGGTGCATGCCTCTA | NA | NA | NA | 0.005802 | 0.005456 |
| C3db | AATGGCCTCCTGAACGTGCGTTGCACTCTTGGGATTTCCTGAGAGTATGTCTGCTTC<br>AGTGCTTAACTTGCCCCAACTTTGCAAGCAGGATGTGTTTCTGCCTTGCATTCTTATG<br>AGCTATTGCCCTCTGAGCCAATGGCTTGTTAATTGCTTGGTTCTTGCAAAATGCTTTG<br>CGCGCTGTTATTCAAGTTTCTACCTTCGTGGTTTTACTTGAGTGACGCTGCTCATGCT<br>TGCAACCGCTGGGATGCAGGTGCATGCCTCTA | NA | NA | NA | 0.005928 | 0.005658 |
| 21209_C | AATGGCCTCCTGAACGTGCGTTGCACTCTTGGGATTTCCTGAAGAGTATGTCTGCTT<br>CAGTGCTTAACTTGCCCCAACTTTGCAAGCAGGATGTGTTTCTGCCTTGCGTTCTTAT<br>GAGCTATTGCCCTCTGAGCCAATGGCTTGTTAATTGCTTGGTTCTTGCAAAATGCTTT<br>GCGCGCTGTTATTCAAGTTTCTACCTTCGTGGTTTTACTTGAGTGACGCTGCTCATGC<br>TTGCAACTGCTGGGATGCAGGTGCATGCCTCTA | NA | NA | NA | 0.009019 | NA |
| 173341_D | AATGGCCCCCTGAACGCGCATTGCACTCTTGGGACTTCCTGAGAGTATGTTTGCTTC<br>AGTGCTTATTTTACCTCCTTGCAAGGTTCTGTCGCAACCTTGTGCCCTGGCCAGCCA<br>TGGGTTAACTTGCCCATGGCTTGCTGAGTAGTGATCTTTTAGAGCAAGCTCTGGCAC<br>GCTGTTGTCTGAGGCAGCCTATATTGAGGCTATTTCAAATGACGTTGCTACAGGCTT<br>GATGTGTCCTTCTGCGCCGTTGCGCATCCCATA | NA | NA | NA | 0.000252 | NA |
| 49135_C | AATGGCCTCCTGAACGTGCGTTGCACTCTTGGGATTTCCTGAGAGTATGTCTGCTTC<br>AGTGCTTAACTTGCCCCAACTTTGCAAGCAGGATGTGTTTCTGCCTTGCGTTCTATG<br>AGCTATTGCCCTCTGAGCCAATGGCTTGTTAATTGCTTGGTTCTTGCAAAATGCTTTG<br>CGCGCTGTTATTCAAGTTTCTACCTTCGTGGTTTTACTTGAGTGACACTGCTCATGCT<br>TGCAACCGCTGGGATGCAGGTGCATGCCTCTA | NA | NA | NA | NA | 0.006062 |
| 40995_C | AATGGCCTCCTGAACGTGCGTTGCACTCTTGGGATTTCCTGAGAGTATGTCTGCTTC<br>AGTGCTTAACTTGCCCCAACTTTGCAAGAAGGATGTGTTTCTGCCTTGCGTTCTTAT<br>GAGCTATTGCCCTCTGAGCCAATGGCTTGTTAATTGCTTGGTTCGTGCAAAATGCTTT<br>GCGCGCTGTTATTCAAGTTTCTACCTTCGTGGTTTTACTTGAGTGACGCTGCTCATGC<br>TTGCAACCGCTGGGATGCAGGTGCATGCCTCTA | NA | NA | NA | NA | 0.007072 |
| C3bd | AATGGCCTCCTGAATGTGCGTTGCACTCTTGGGATTTCCTGAGAGTATGTCTGCTTC<br>AGTGCTTAACTTGCCCCAACTTTGCAAGCAGGATGTGTTCCTGCCTTGCGTTCGTAT<br>GAGCTGTTGCCCTCTGAGCCAATGGCTTGTTAATTGCTTGGTTCTTGCAAAATGCTTT<br>GCGCGCTGTTATTCAAGTTTCTACCTTCGTGGTTTTACTTGAGTGATGCTGCTCATGC<br>TTGCAACCGCTGGGATGCAGGTGCATGCCTCTA | NA | NA | NA | NA | 0.006567 |

**Table S5.** Sampling summary, library statistics, and symbiont composition of *Hydnophora microconos* across intra-colony sample positions (1-5: apex-base).

| Library statistic | Position 1 | Position 2 | Position 3 | Position 4 | Position 5 |
|---|---|---|---|---|---|
| Total number of sequences | 847480 | 962695 | 836588 | 910665 | 930736 |
| Total number of samples | 49 | 49 | 46 | 49 | 48 |
| Average sequence depth per sample (+/-SD) | 17296 (+/-6401) | 19647 (+/-6648) | 18187 (+/-6049) | 18585 (+/-5290) | 19390 (+/-5310) |
| Proportion profile sequences | 0.76 | 0.78 | 0.76 | 0.76 | 0.80 |
| Proportion non-profile sequences | 0.24 | 0.22 | 0.24 | 0.24 | 0.20 |
| Total number ITS2 profiles | 17 | 9 | 10 | 9 | 13 |
| Proportion of samples with >1 ITS2 profiles | 0.45 | 0.43 | 0.46 | 0.37 | 0.60 |
| Number of C3k\|D associated samples | 11 | 11 | 9 | 9 | 12 |
| Number of C3f\|D associated samples | 7 | 10 | 11 | 8 | 13 |

A)

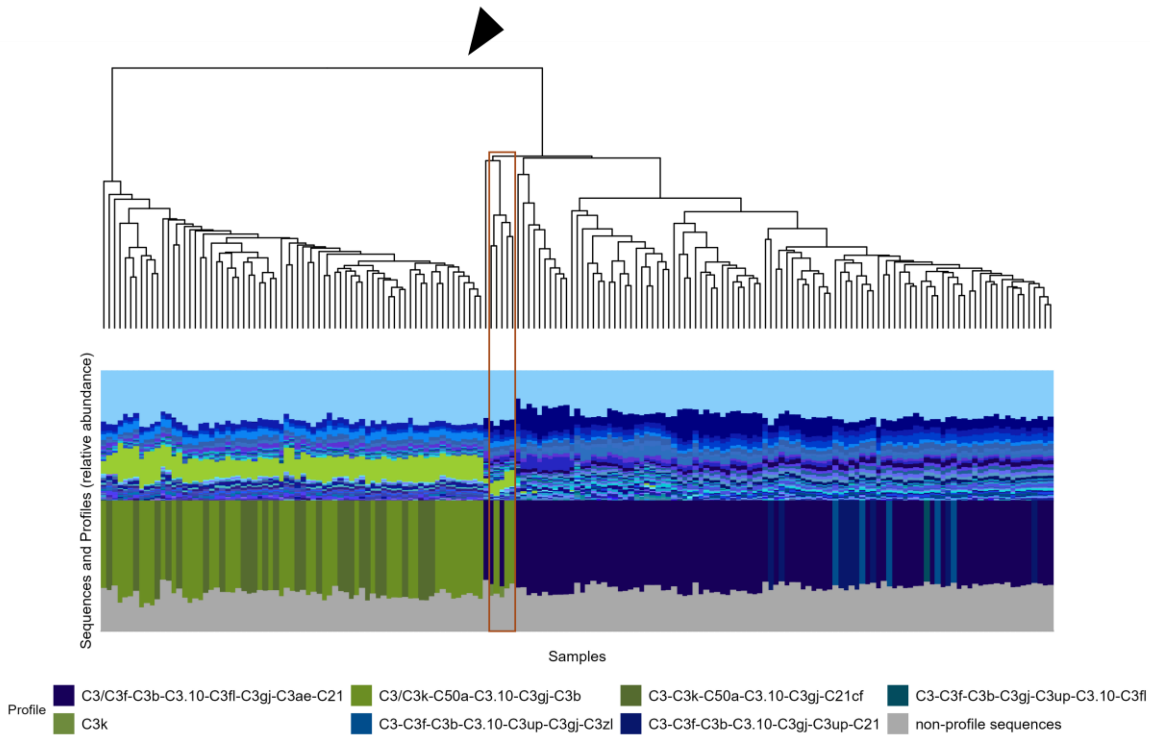


B)

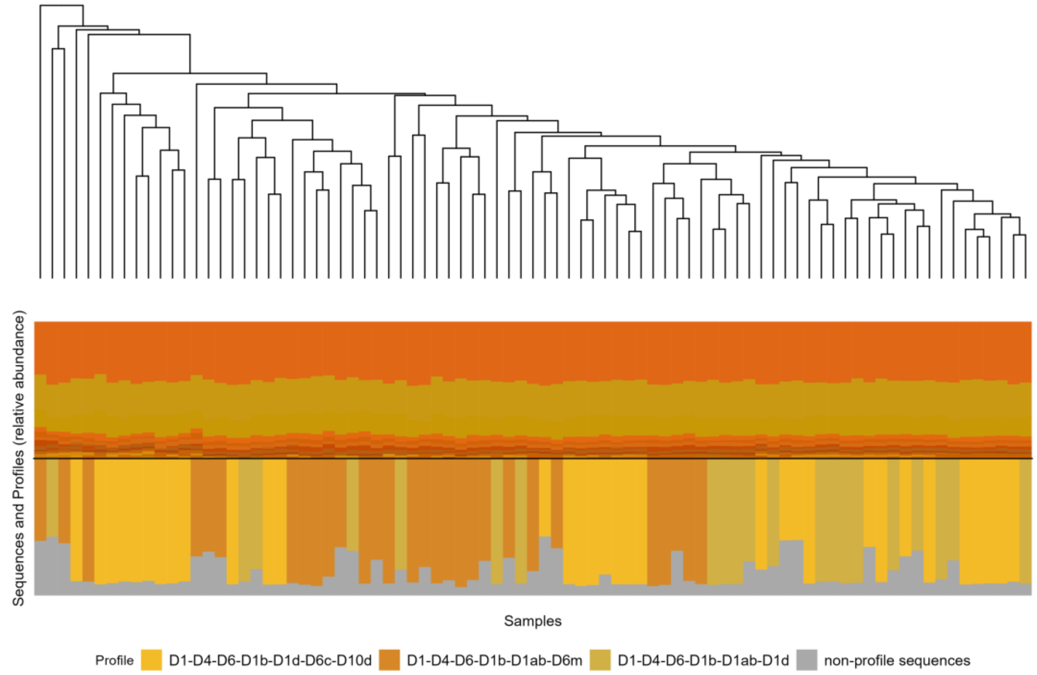


**Figure S1. A)** K-mer-based hierarchical clustering of per-sample *Cladocopium* sequences (top) together with ITS2 sequence (middle) and profile (base) relative abundance. Black triangle indicates major split into C3k (left) and C3f (right) lineages. Red box indicates five samples where intermediate contributions of C3f and C3k were detected (see *Linear deconvolution* above) **B)** K-mer-based hierarchical clustering of per-sample *Durusdinium* sequences (top) together with ITS2 sequence (middle) and profile (base) relative abundance.

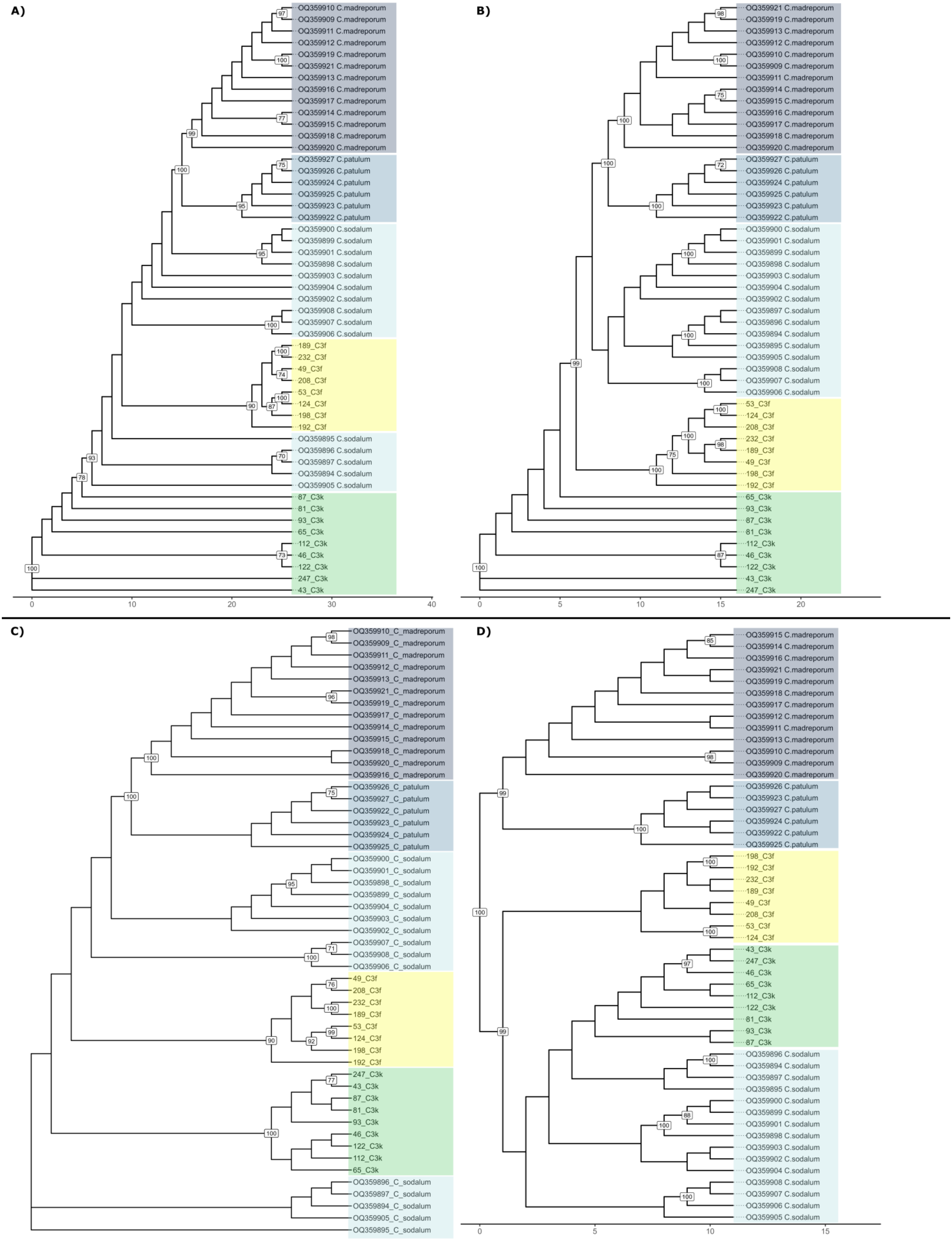

A)
OQ359910 C.madreporum
OQ359909 C.madreporum
OQ359911 C.madreporum
OQ359912 C.madreporum
OQ359919 C.madreporum
OQ359921 C.madreporum
OQ359913 C.madreporum
OQ359916 C.madreporum
OQ359917 C.madreporum
OQ359914 C.madreporum
OQ359915 C.madreporum
OQ359918 C.madreporum
OQ359920 C.madreporum
OQ359927 C.patulum
OQ359926 C.patulum
OQ359924 C.patulum
OQ359925 C.patulum
OQ359923 C.patulum
OQ359922 C.patulum
OQ359900 C.sodalum
OQ359899 C.sodalum
OQ359901 C.sodalum
OQ359898 C.sodalum
OQ359903 C.sodalum
OQ359904 C.sodalum
OQ359902 C.sodalum
OQ359908 C.sodalum
OQ359907 C.sodalum
OQ359906 C.sodalum
189_C3f
232_C3f
49_C3f
208_C3f
53_C3f
124_C3f
198_C3f
192_C3f
OQ359895 C.sodalum
OQ359896 C.sodalum
OQ359897 C.sodalum
OQ359894 C.sodalum
OQ359905 C.sodalum
87_C3k
81_C3k
93_C3k
65_C3k
112_C3k
46_C3k
122_C3k
247_C3k
43_C3k
0
10
20
30
40
B)
OQ359921 C.madreporum
OQ359919 C.madreporum
OQ359913 C.madreporum
OQ359912 C.madreporum
OQ359910 C.madreporum
OQ359909 C.madreporum
OQ359911 C.madreporum
OQ359914 C.madreporum
OQ359915 C.madreporum
OQ359916 C.madreporum
OQ359917 C.madreporum
OQ359918 C.madreporum
OQ359920 C.madreporum
OQ359927 C.patulum
OQ359926 C.patulum
OQ359924 C.patulum
OQ359925 C.patulum
OQ359923 C.patulum
OQ359922 C.patulum
OQ359900 C.sodalum
OQ359901 C.sodalum
OQ359899 C.sodalum
OQ359898 C.sodalum
OQ359903 C.sodalum
OQ359904 C.sodalum
OQ359902 C.sodalum
OQ359897 C.sodalum
OQ359896 C.sodalum
OQ359894 C.sodalum
OQ359895 C.sodalum
OQ359905 C.sodalum
OQ359908 C.sodalum
OQ359907 C.sodalum
OQ359906 C.sodalum
53_C3f
124_C3f
208_C3f
232_C3f
189_C3f
49_C3f
198_C3f
192_C3f
65_C3k
93_C3k
87_C3k
81_C3k
112_C3k
46_C3k
122_C3k
43_C3k
247_C3k
0
5
10
15
20
C)
OQ359910_C_madreporum
OQ359909_C_madreporum
OQ359911_C_madreporum
OQ359912_C_madreporum
OQ359913_C_madreporum
OQ359921_C_madreporum
OQ359919_C_madreporum
OQ359917_C_madreporum
OQ359914_C_madreporum
OQ359915_C_madreporum
OQ359918_C_madreporum
OQ359920_C_madreporum
OQ359916_C_madreporum
OQ359926_C_patulum
OQ359927_C_patulum
OQ359922_C_patulum
OQ359923_C_patulum
OQ359924_C_patulum
OQ359925_C_patulum
OQ359900_C_sodalum
OQ359901_C_sodalum
OQ359898_C_sodalum
OQ359899_C_sodalum
OQ359904_C_sodalum
OQ359903_C_sodalum
OQ359902_C_sodalum
OQ359907_C_sodalum
OQ359908_C_sodalum
OQ359906_C_sodalum
49_C3f
208_C3f
232_C3f
189_C3f
53_C3f
124_C3f
198_C3f
192_C3f
247_C3k
43_C3k
87_C3k
81_C3k
93_C3k
46_C3k
122_C3k
112_C3k
65_C3k
OQ359896_C_sodalum
OQ359897_C_sodalum
OQ359894_C_sodalum
OQ359905_C_sodalum
OQ359895_C_sodalum
D)
OQ359915 C.madreporum
OQ359914 C.madreporum
OQ359916 C.madreporum
OQ359921 C.madreporum
OQ359919 C.madreporum
OQ359918 C.madreporum
OQ359917 C.madreporum
OQ359912 C.madreporum
OQ359911 C.madreporum
OQ359913 C.madreporum
OQ359910 C.madreporum
OQ359909 C.madreporum
OQ359920 C.madreporum
OQ359926 C.patulum
OQ359923 C.patulum
OQ359927 C.patulum
OQ359924 C.patulum
OQ359922 C.patulum
OQ359925 C.patulum
198_C3f
192_C3f
232_C3f
189_C3f
49_C3f
208_C3f
53_C3f
124_C3f
43_C3k
247_C3k
46_C3k
65_C3k
112_C3k
122_C3k
81_C3k
93_C3k
87_C3k
OQ359896 C.sodalum
OQ359894 C.sodalum
OQ359897 C.sodalum
OQ359895 C.sodalum
OQ359900 C.sodalum
OQ359899 C.sodalum
OQ359901 C.sodalum
OQ359898 C.sodalum
OQ359903 C.sodalum
OQ359902 C.sodalum
OQ359904 C.sodalum
OQ359908 C.sodalum
OQ359907 C.sodalum
OQ359906 C.sodalum
OQ359905 C.sodalum
0
5
10
15

**Figure S2.** The position of psbA[ncr] samples relative to recent *Cladocopium* description [7], labelled by ITS2 *Cladocopium* C3f or C3k identification across four tree-building methods; **A)** maximum parsimony **B)** maximum parsimony with indel as 5[th] character state **C)** maximum likelihood **D)** alignment-free k-mer with jack-knifing.

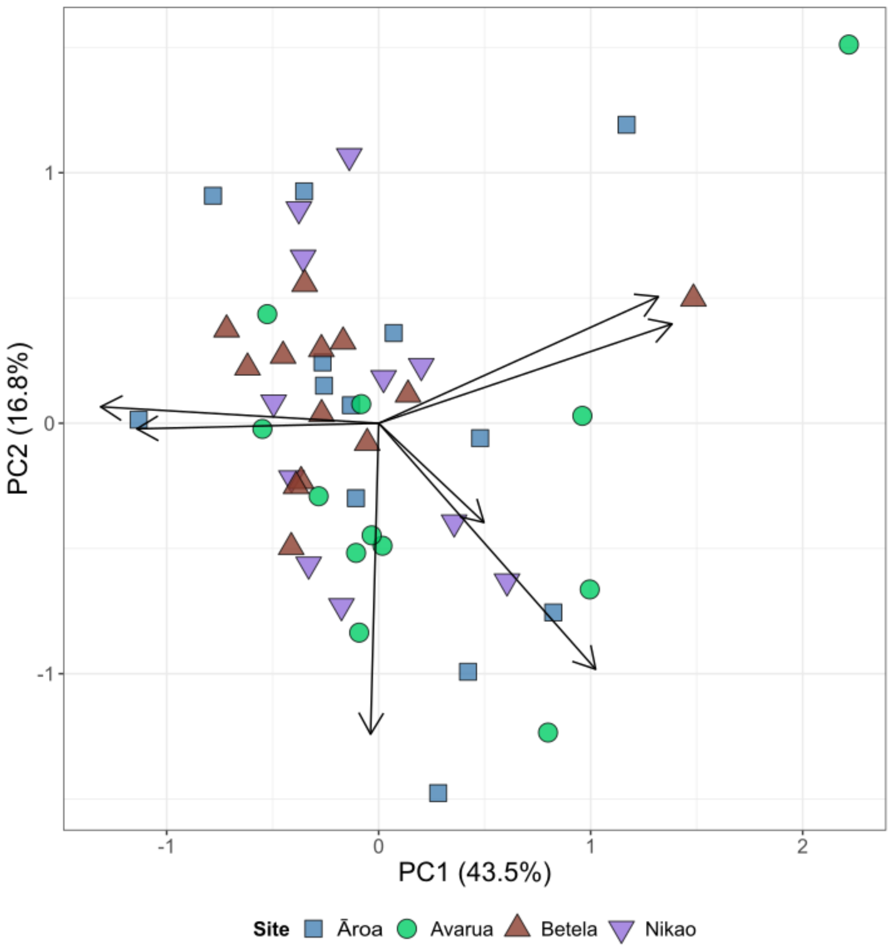


**Figure S3.** Principal Components Analysis of colony morphometrics. Each point represents one *Hydnophora microconos* colony. Points are coloured and shaped according to sampling site.

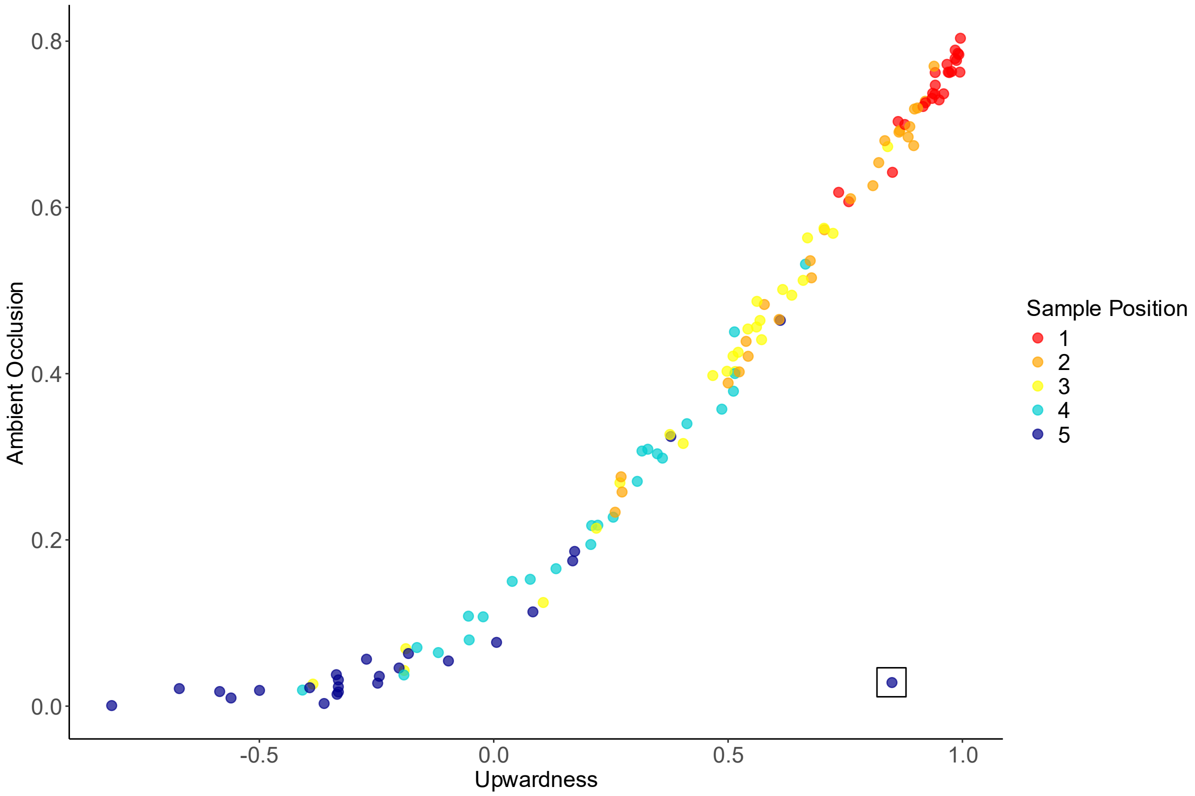


**Figure S4.** Relationship between upwardness and ambient occlusion. Each point represents an individual sample, coloured by sample position. Black box indicates single upward oriented, highly occluded, base region outlier sample.

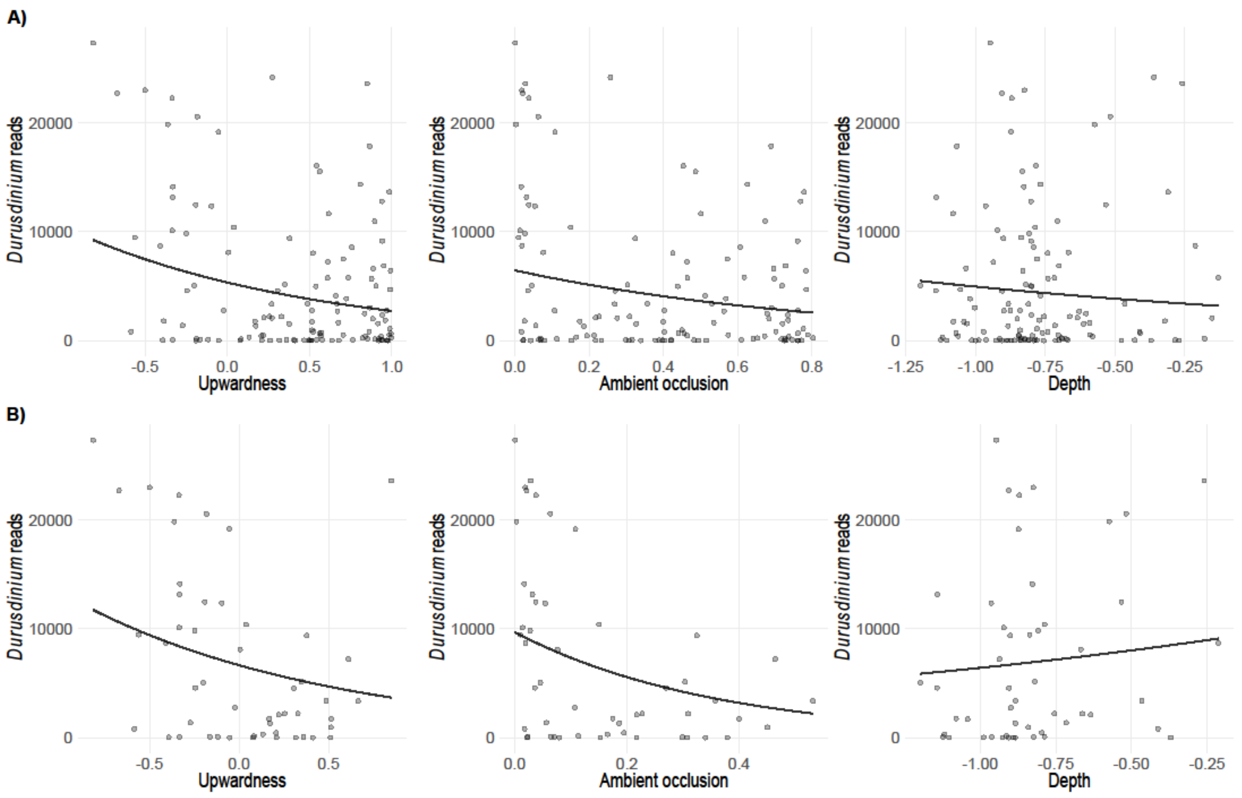


**Figure S5. A)** Zero-inflated negative binomial generalised linear mixed models for the variables 'upwardness', 'ambient occlusion', and 'depth', predicting *Durusdinium* read count across all intra-colony samples (1-5) for colonies with multi-symbiont assemblages. **B)** Zero-inflated negative binomial generalised linear mixed models for the variables 'upwardness', 'ambient occlusion', and 'depth', predicting *Durusdinium* read count across the colony base region (sample positions 4 and 5) for colonies with multi-symbiont assemblages, and outlier sample (see Figure S4) included. All models included colony-specific zero-inflation terms and colony as a random intercept. Points represent individual samples; the black line represents the model-predicted reads. Models excluded the site Avarua.

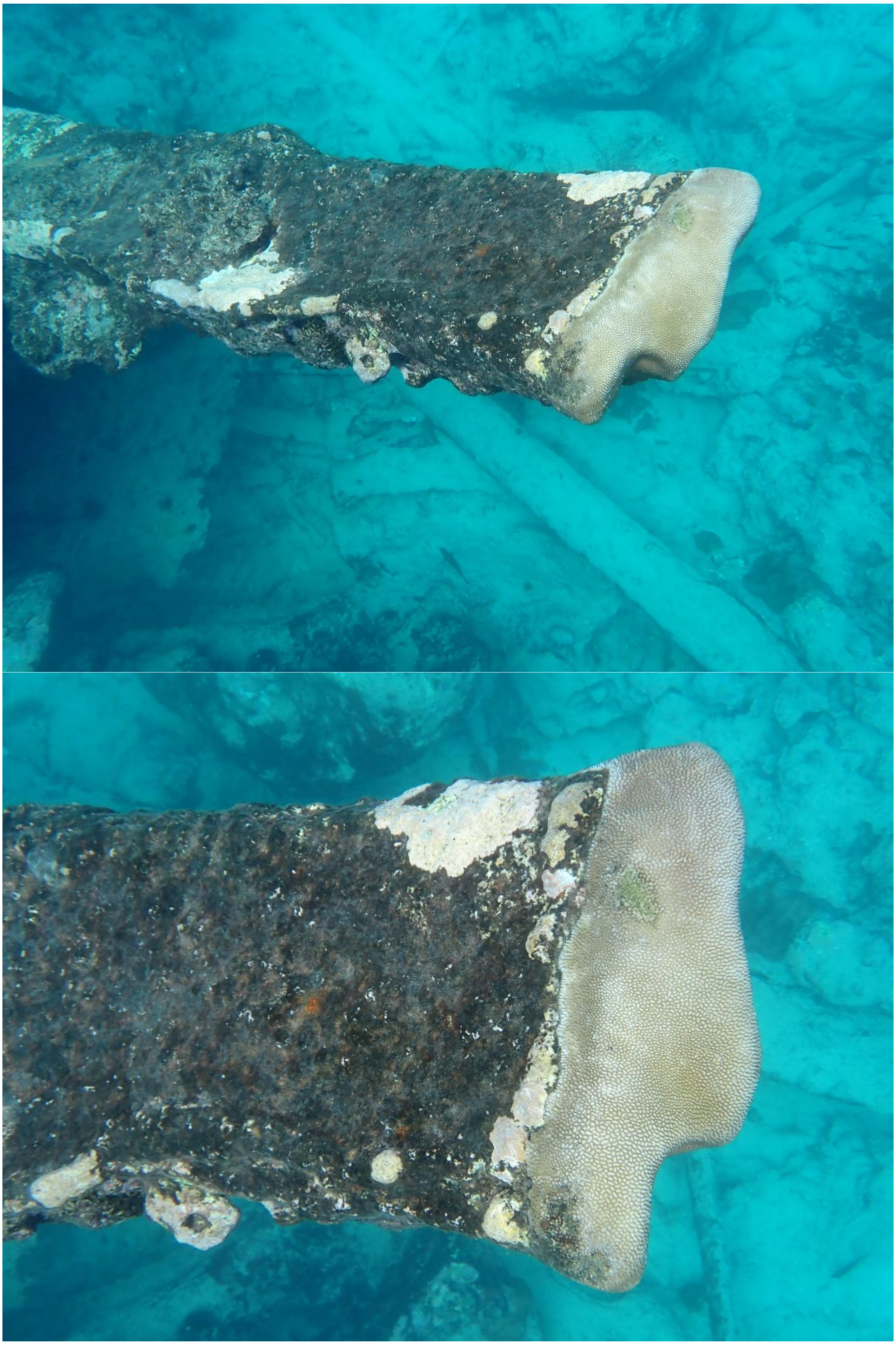

**Figure S6.** Example images of *Hydnophora microconos* exhibiting encrusting growth form, taking on the morphology of the supporting structure. Photo credit Phoebe Argyle
